# Serotonin selectively modulates visual responses of object motion detectors in *Drosophila*

**DOI:** 10.1101/2025.03.21.644681

**Authors:** David J. Bertsch, Lesly M. Palacios Castillo, Mark A. Frye

## Abstract

Serotonin (5-HT) is a hormonal messenger that confers state-level changes upon the nervous system in both humans and flies. In *Drosophila*, lobula columnar (LC) cells are feature-detecting neurons that project from the optic lobe to the central brain, where each population forms an anatomically-distinct glomerulus with heterogeneous synaptic partners. Here, we investigated serotonin’s effect on two LC populations with different 5-HT receptor expression profiles. Receptor expression does not predict neuromodulatory effects: LC15 expresses inhibitory 5-HT1A and 5-HT1B receptors, yet serotonin increases the amplitude of calcium responses to visual stimuli. LC12 expresses inhibitory 5-HT1A and excitatory 5-HT2A receptors, yet serotonin application does not influence visual responses. Serotonin targets select visual response properties, potentiating LC15 responses to a motion-defined bar and tuning the gain of responses to varying object velocity but has no influence on contrast sensitivity. Serotonin does not significantly facilitate LC15 responses in postsynaptic dendrites, only in the presynaptic terminals of the glomerulus, which suggests that the neuromodulatory effects are strongest in the central brain. Connectomics confirms that LC12 and LC15 share neither presynaptic inputs nor postsynaptic outputs in the central brain. The wiring diagram shows no synaptic interactions between the LC15 circuit and major serotonergic 5-HTPLP neurons, nor to other serotonergic neurons of the central brain, suggesting that endogenous 5-HT acts via paracrine transmission on non-serotonergic pathways. Lobula- and glomerulus-specific GABAergic and glutamatergic inhibitory partners, positioned to filter visual stimuli, are putative 5-HT targets. These results provide a comparative framework for the neuromodulatory mechanisms involved in visual processing.

**New & Noteworthy:** How does neuromodulatory state affect visual feature detection? In this work, we demonstrate highly specific facilitated visual responses of object-detecting neurons after serotonin bath application in Drosophila. Serotonin potentiates motion-defined bar responses in object-detecting LC15 neurons and tunes response gain to translating bars of mid-range velocities in presynaptic axon terminals. Our calcium imaging extends what is known about extra-synaptic neuromodulation in the visual system and shows that serotonin heightens visual processes that inform object-specific behavior.

## Introduction

Visual animals must extract object information from a complex, changing external environment while receiving internal physiological cues to produce context-specific behavior. Endogenous release of signaling molecules communicates state-dependent changes in all animals yet we understand neither how these neuromodulators transform visual information as it enters the central brain nor how they reconfigure the hard-wired and numerically-limited circuits of the brain to create flexible state-dependent behavior (1). An important class of neuromodulatory chemicals is the biogenic monoamines which impart remarkable flexibility to both vertebrate and invertebrate neural networks (2). The biogenic amines activate G protein-coupled receptors (GPCRs) on target cells and include dopamine (DA), serotonin (5-HT), tyramine (TA), norepinephrine (NE) and the invertebrate analog, octopamine (OA) (3–6). Serotonin (5-hydroxytryptamine, 5-HT) is a pleiotropic neurotransmitter and neurohormone that mediates learning and memory, sleep, feeding, aggression, arousal, courtship, swarming behavior in locusts, and locomotion as well as various developmental or physiological processes including critical period plasticity in both vertebrates and invertebrates (7–17). Importantly, 5-HT has been implicated in enhancing gain control in the glomeruli of the olfactory antennal lobe (18–20), and for enhanced visual responses by lamina class L2 in response to OFF objects (i.e. objects darker than the background) (21).

The genetically tractable fly *Drosophila melanogaster* has been used for decades to understand visual processing and neuromodulatory control of behavior. Detailed reviews have summarized how the anatomy and physiology of the optic lobe relates to visual information processing (22–25). In brief, luminance changes in the environment are detected by photoreceptors in the retina (R1-R6) before being transmitted downstream to the optic lobe. The optic lobe is composed of four neuropils: the lamina, medulla, lobula, and lobula plate. Each neuropil shows anatomically distinct ‘horizontal’ processing layers and retinotopic arrays of ‘vertical’ columns where luminance signals are processed for directional motion, features, or color. Columnar signals are spatially integrated by visual projection neurons (VPNs) that send axons to higher brain areas. The innermost neuropil of the optic lobe, the lobula (Lo), houses a class of VPNs called lobula columnar (LC) cells. The LCs include 22 neural populations that encode select visual features of relevant objects in the fly’s environment (26,27). Each LC type is made up of numerous clones of similar morphology that tile the visual field. Dendrites integrate synaptic inputs across a set of visual columns, the number of which varies for each LC type, and project axons to the central brain where the terminals of all members of a given type converge with postsynaptic neurons within an anatomically-distinct optic glomerulus (OG) in the anterior or posterior ventrolateral protocerebrum (AVLP, PVLP) (28–31). LC12 and LC15 both respond to translating bars, which are highly engaging to a flying fly as they represent landscape features moving against the visual surroundings (32). Bars may be defined by luminance (darker or brighter than the visual surroundings) or by optical disparities generated by translational cues relative to the stationary ground (i.e. motion). Neither LC12 nor LC15 respond to wide-field motion (square grating period 17.6°), and both types have their postsynaptic dendrites in Lo layers 2 and 4. It has been demonstrated using two-photon calcium imaging that LC cells can be measured individually in the Lo whereas the population of axon terminals can be measured collectively at the terminal OG (33,34). Within the OG, LC12 and LC15 axon terminals make excitatory and inhibitory synapses with other cell types. We have previously shown that both LC12 and LC15 show modulation by bath application of an octopamine agonist (35). LC responses to local features have previously been measured using single-cell or pan-neuronal imaging and functionally grouped (26,36). In one report, LC15 was grouped by strong responses to moving objects, with weak responses to looming objects, whereas LC12 was grouped by its strong responses to looming and preference for OFF over ON contrast polarity (26). A second study placed both LC12 and LC15 in the same group defined by strong responses to vertical bars as well as medium and large moving spots, plus some looming sensitivity (36). Disparities in LC response classification could be due to the specific approach used, or by higher order neuromodulation of complex feature detectors.

We explored the effects of 5-HT on LC responses in the optic lobe and optic glomeruli of the central brain of *Drosophila melanogaster* using pharmacology, single-cell intersectional genetics, connectomics, and two-photon calcium imaging. To mimic the effects of volumetric neurohormonal release, we bath-applied serotonin HCl in *Drosophila* saline while performing two-photon calcium imaging. Imaging revealed that 5-HT bath application enhances LC15 responses to motion-defined vertical bars. We observed that the lobular dendrites showed non-significant but increased Ca^2+^ response strength, whereas the glomerular axon terminals in the PVLP showed significant potentiation after 5-HT. We next presented OFF bars moving at different speeds and observed that 5-HT significantly increases the LC15 glomerular response to mid-range velocities. To determine if this serotonergic potentiation is a general phenomenon among bar-sensitive feature detectors, we showed the same bar stimuli while imaging the LC12 glomerulus and did not observe any modulatory change. Using the synaptic connectome (37), we detail LC-specific GABAergic inhibitory neurons that display reciprocal innervation of the LCs and could mediate glomerular potentiation. Both VPN types respond to translating bar stimuli but only LC15 shows potentiated Ca^2+^ responses to specific objects after 5-HT bath application. Our data support the hypothesis that serotonergic potentiation occurs within axon terminals of the VLP and the two LC types respond to the neuropeptide in ways that are likely determined by their specific synaptic partners rather than their 5-HTR expression profile.

## Materials and Methods

### Fly stocks

*Drosophila melanogaster* were maintained on a standard cornmeal and molasses-based agar medium with a 12:12 hour light/dark cycle at room temperature (22-25°C) in 40 ml vials. Only 3-5 day old adult females were used for experimentation. To genetically target LC neurons we exclusively used split-GAL4 driver lines because of their high specificity for the respective class of neurons. OL0042B was the driver line used to target LC15 and OL0007B to target LC12 (27). These flies were crossed with virgin females expressing UAS-GCaMP6f and the progeny were used for experiments (38).

### Animal preparation

Imaging experiments were performed between ZT0–14, although time of day was not a factor in our experimental design or analysis. Female flies were anesthetized at 4°C on a cold plate and mounted to a custom 3D printed fly holder using ultraviolet glue (Form3 SLA printer, FormLabs; Dreve Fotoplast gel, Audiology Supplies, 44811). The fly’s legs were immobilized with low melting point beeswax to eliminate interference with recordings and visual stimulation. Fine forceps (Dumont, #5SF, Fine Science Tools) were used to remove the cuticle on the posterior surface of the fly’s head to expose the right optic lobe in the region of the lobula and VLP. We severed muscles 1 and 16 to reduce brain movements. This approach has previously been used in our laboratory (35,39).

### Pharmacology and bath perfusion

*D. melanogaster* saline was composed of ultrapure water (18.3 MU, Millipore) and (in mM): 103 NaCl, 3 KCl, 1.5 CaCl_2_, 4 MgCl_2_, 26 NaHCO_3_, 1 NaH_2_PO_4_, 10 trehalose, 10 glucose, 5 TES, 2 sucrose, pH 7.0 - 7.25. We tested the pH value of the final mixed solutions using an Orion Star A211 pH meter (Thermo Scientific) and did not observe any changes outside of the normal range at room temperature. pH drift was not observed in refrigerated saline until 6 days after at this TES concentration. Serotonin HCl (Sigma Aldrich, cat# H9523) stock solutions were stored at 20°C in small quantities and diluted in *D. melanogaster* saline to the final concentration of 100 μM. All 5-HT solutions were kept away from light. All perfusate solutions were oxygenated during bath perfusion (95% O^2^, 5% CO^2^) using a saltwater aquarium airstone (Alegi). Measurements were taken after a 10-minute wash-in. During two-photon imaging, the brain was continuously perfused with freshly prepared extracellular saline or 5-HT HCl solution at 1.5 ml/min via a computer-controlled six channel valve controller (VC-6, Warner Instruments) and a custom-built gravity-drip perfusion system. Bath temperature was kept at 20°C with an inline-solution heater (SC-20, Warner Instruments) and a temperature controller (TC-324, Warner Instruments).

### Two-photon calcium imaging

LC neurons were imaged at 920 nm using a Ti:Sapphire pulse laser (Chameleon Vision, Coherent, Santa Clara, CA) controlled by SlideBook (Version 6, 3i, Boulder, CO. RRID:SCR_014300). We imaged with a 20x water-immersion objective (W Plan-Apochromat, 1.0 DIC, Zeiss) with three layers of blue filter (Indigo, Rosco, No. 59) to reduce bleed-through from the LED arena to the photomultipliers. Single plane images were taken at 7.0-10.0 frames/s with each frame at a x-y pixel resolution ranging from 150 x 256 to 168 x 212 and 0.2 to 0.3 mm pixel spacing. Images and external stimulations were synchronized *a posteriori* using frame capture markers (TTL pulses output from Slidebook) and stimulus events (analog outputs from the LED display controller) sampled with a data acquisition device (DAQ) (PXI-6259, NI) at 10 kHz. The DAQ interfaced with MATLAB (R2020a, MathWorks. RRID:SCR_001622) via rack-mount terminal block (BNC-2090, NI). To record population responses of a given LC type, GCaMP6f responses were recorded in the LC output glomeruli where all terminals merge together. To record activity of single LC neurons, GCaMP6f responses were collected at different z-planes from individual dendrites in the lobula.

### Visual stimulation

Visual stimuli during two-photon imaging were presented on a cylindrical LED arena (40). The arena was composed of 4 rows and 12 columns of 8 x 8 LED dot matrix panels (470nm, Dongguan Houke Electronic, 12088-AB). The LED display covered ±108° of the visual field in the horizontal, and ±35° in the vertical dimension. The arena subtends 216° in azimuth by 70° in elevation, comprising a dot matrix array of 96 x 32 pixels. Each pixel is 3.0mm in diameter separated by 3.97mm center-to-center. Each pixel subtended to 2.2° on the fly’s retina at the visual equator. The display had a maximum intensity of ∼0.11 μW m^−2^ (recorded at the fly’s position) at the spectral peak of 460 nm (full width at half maximum: 243 nm). The ‘motion-defined’ bar stimulus (Figure 1, Figure 2) had an average irradiance of ∼0.06 μW m^−2^. Stimuli were generated and controlled using custom Matlab scripts. The mean irradiance values of the luminance series bar stimuli (Figure 3) were measured as 0.0, 0.009, 0.018, 0.039, 0.074, 0.092, 0.101, and 0.11 μW m^−2^. In Figure 3Bii, Weber contrast is calculated by subtracting the light intensity of the object from the background and dividing by the background intensity.

**Figure 1.**
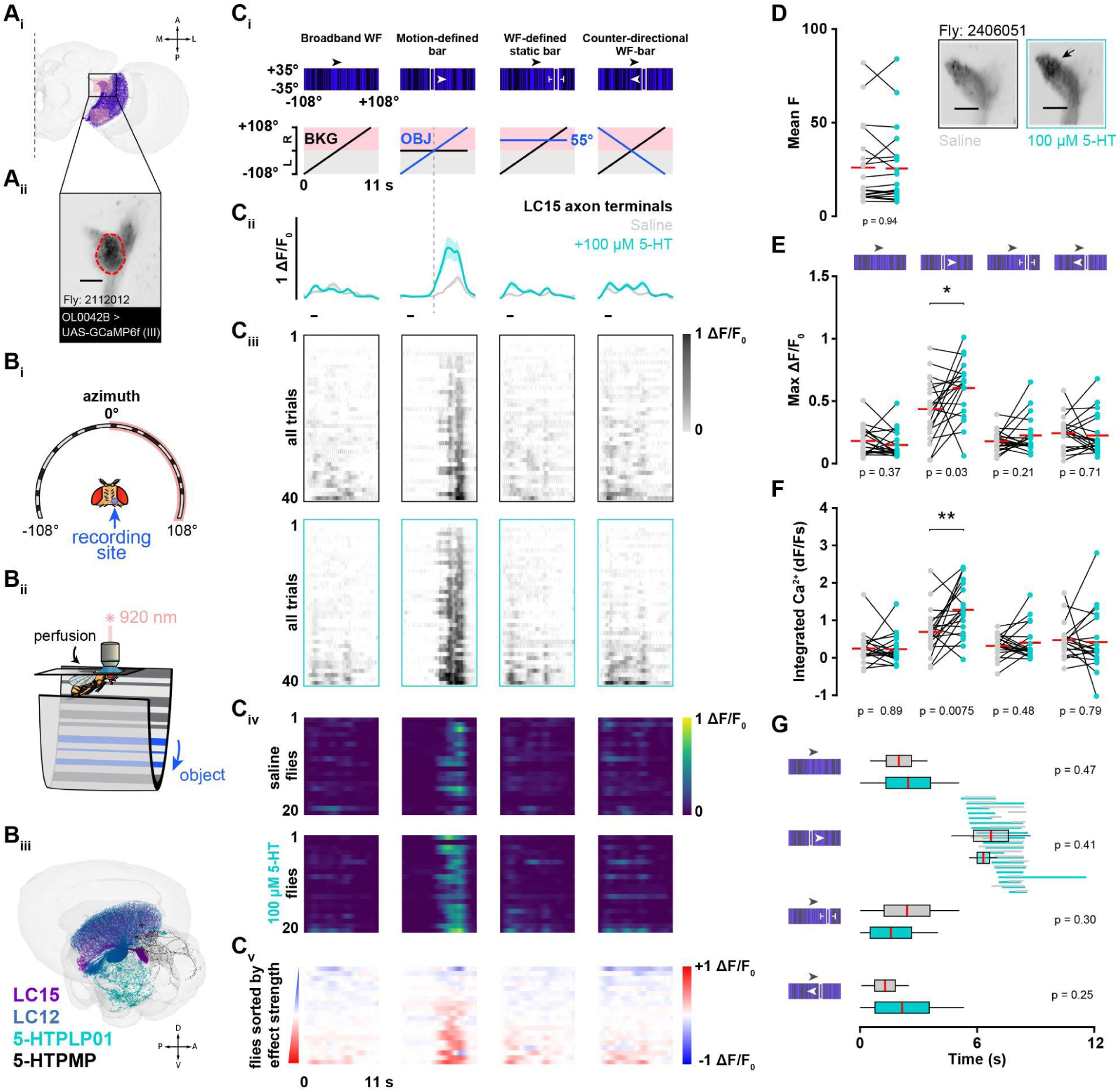
The LC15 glomerular response to a motion-defined bar is potentiated by 5-HT when the background is stationary. (Ai) LC15 cells (purple) in the right hemisphere. The two pink boxes highlight the recording sites: glomerulus of population axon terminals (top box) and individual dendritic projections (bottom box). (Aii) Two-photon fluorescent image of z-axis projection showing the glomerulus of LC15 cells. The red dotted line defines the region of interest. Scale bar, 25 µm. (Bi) Cartoon of head-fixed preparation presented with visual stimuli from below using a wrap-around LED display that extends +/- 108° in azimuth. All recordings were collected from the right hemisphere. Not drawn to scale. (Bii) Schematic showing the two-photon imaging preparation, with 20x water immersion lens. Temperature-controlled *Drosophila* saline was perfused into the preparation for the duration of the experiment. (Biii) LC15 (purple) and LC12 (blue) do not directly synapse with 5-HTPLP neurons (cyan) in the right hemisphere but likely receive 5-HT through paracrine release into the AVLP and PVLP. Projections from 5-HTPMP are shown in black. Reconstruction generated using FlyWire.ai (37). (Ci) Schematic illustrating the visual stimulus presented on the LED display (top) and the motion trajectory of object (OBJ) or background (BKG) (bottom). (Cii) Mean GCaMP6f signal from the LC15 axon terminals (±SEM shading) before (gray) and after 100 µM 5-HT (cyan). The horizontal scale bar (black) represents 1 second. (Ciii) ΔF/F_0_ response raster plots displaying individual trials before and after 100 µM 5-HT (cyan). Two repeats of each stimulus were shown to each animal (N = 20, n = 40). The trial responses are sorted by increasing peak ΔF/F_0_ response. (Civ) ΔF/F_0_ response raster plots displaying mean animal responses before (top) and after 100 µM 5-HT (bottom) (N = 20). Flies 1-20 (rows) are sorted the same in the pre-5-HT (top) and post-5-HT (bottom) condition rasters. (Cv) Mean change in effect size rasters of LC15 glomerular GCaMP6f responses for each animal in the saline and 5-HT conditions. Animals (row) are sorted by increasing effect size strength. (D) Pairwise comparison of mean fluorescent intensity in the LC15 glomerulus before (gray) and after 100 µM 5-HT (cyan). Horizontal red lines correspond to the population means. Two-tailed paired t-test, N = 20. Insert: average frames of LC15 fluorescence intensity from a single animal before (gray) and after 100 µM 5-HT (cyan). Arrow points to the distal tip of the LC15 glomerulus where the presynaptic terminals are located. Scale bar, 25 µm. (E) Pairwise comparison of maximum ΔF/F_0_ responses in the LC15 glomerulus before (gray) and after 100 µM 5-HT (cyan). Two-tailed paired t-test, N = 20. (F) Pairwise comparison of integrated Ca^2+^ responses in the LC15 glomerulus before (gray) and after 100 µM 5-HT (cyan). Two-tailed paired t-test, N = 20. (G) Pairwise comparison of time to half-max (s) for LC15 glomerular responses before (gray) and after 100 µM 5-HT (cyan). Two-tailed paired t-test, N = 20. The horizontal lines behind the box plots in the motion-defined bar condition indicate the time period from the half-max to the maximum ΔF/F_0_ responses for each animal before (gray) and after 100 µM 5-HT (cyan).

**Figure 2.**
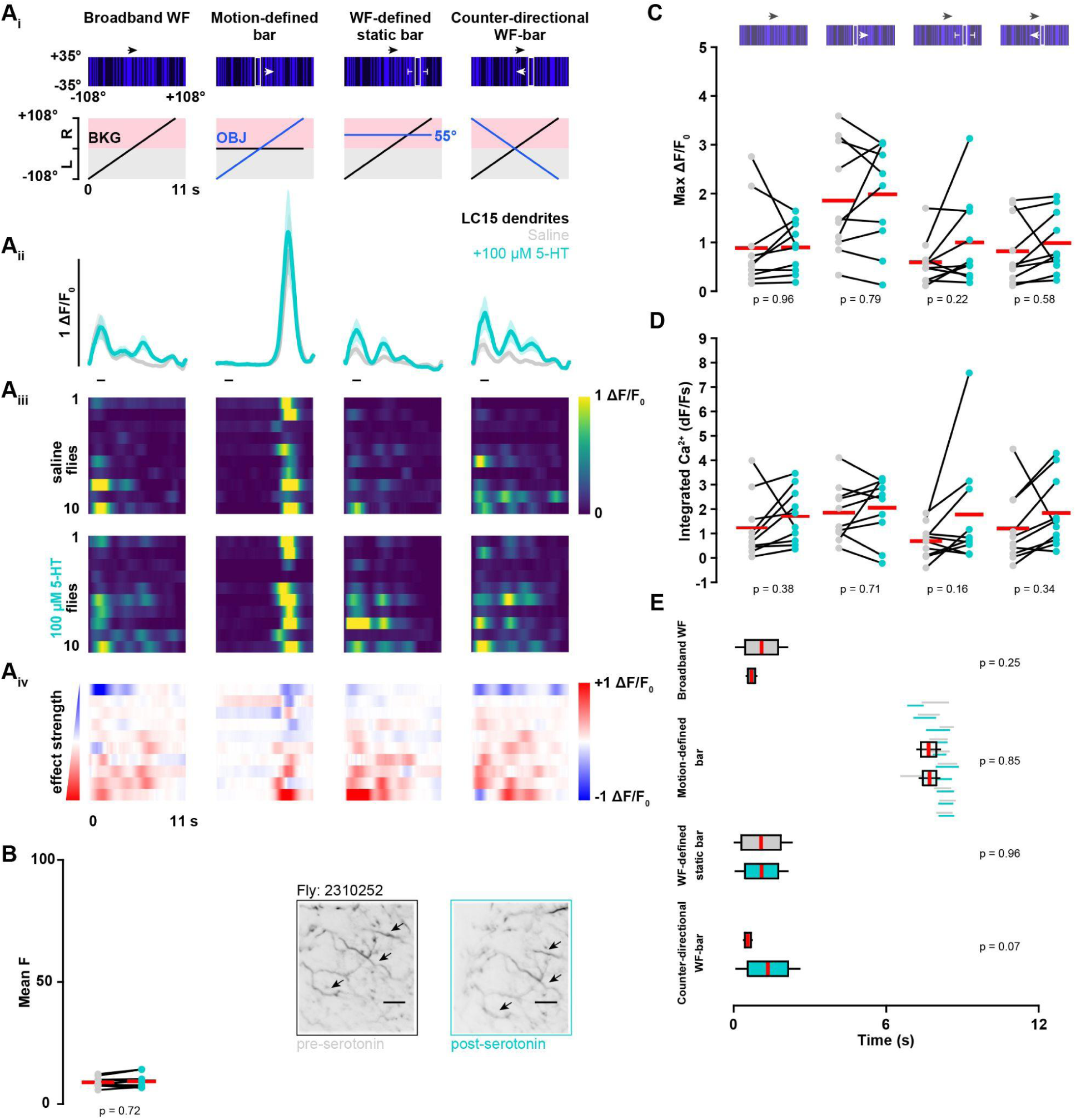
LC15 dendrites do not show potentiated responses after 5-HT bath perfusion. (Ai) Schematic illustrating the visual stimulus presented on the LED display (top) and the motion of the visual feature elements over time (bottom). (Aii) Population mean GCaMP6f signal from the LC15 dendrites (±SEM shading) before (gray) and after 100 µM 5-HT (cyan). The vertical scale bar (black) represents 1 ΔF/F_0_. The horizontal scale bar (black) represents 1 second. (Aiii) ΔF/F_0_ response raster plots displaying mean animal responses before (top) and after 100 µM 5-HT (bottom) (N = 10). Flies 1-10 (rows) are sorted the same in the pre-5-HT (top) and post-5-HT (bottom) condition rasters. (Aiv) Mean change in effect size rasters of LC15 dendritic GCaMP6f responses for each animal. Animals (row) are sorted by increasing effect size strength. (B) Pairwise comparison of mean fluorescent intensity in the LC15 dendrites before (gray) and after 100 µM 5-HT (cyan). Horizontal red lines correspond to the population means. Two-tailed paired t-test. Mean time lapse between pre- and post-conditions = +38 minutes. Insert: average frames of LC15 fluorescence intensity from a single animal before (gray) and after 100 µM 5-HT (cyan). Scale bar, 25 µm. (C) Pairwise comparison of maximum ΔF/F_0_ responses in the LC15 dendrites before (gray) and after 100 µM 5-HT (cyan). Two-tailed paired t-test, N = 10. (D) Pairwise comparison of integrated Ca^2+^ responses in the LC15 dendrites before (gray) and after 100 µM 5-HT (cyan). Two-tailed paired t-test, N = 10. (E) Pairwise comparison of time to half-max (s) for LC15 dendrite responses before (gray) and after 100 µM 5-HT (cyan). Two-tailed paired t-test, N = 10. The horizontal lines behind the box plots in the motion-defined bar condition indicate the time period from the half-max to the maximum ΔF/F_0_ responses for each animal before (gray) and after 100 µM 5-HT (cyan).

**Figure 3.**
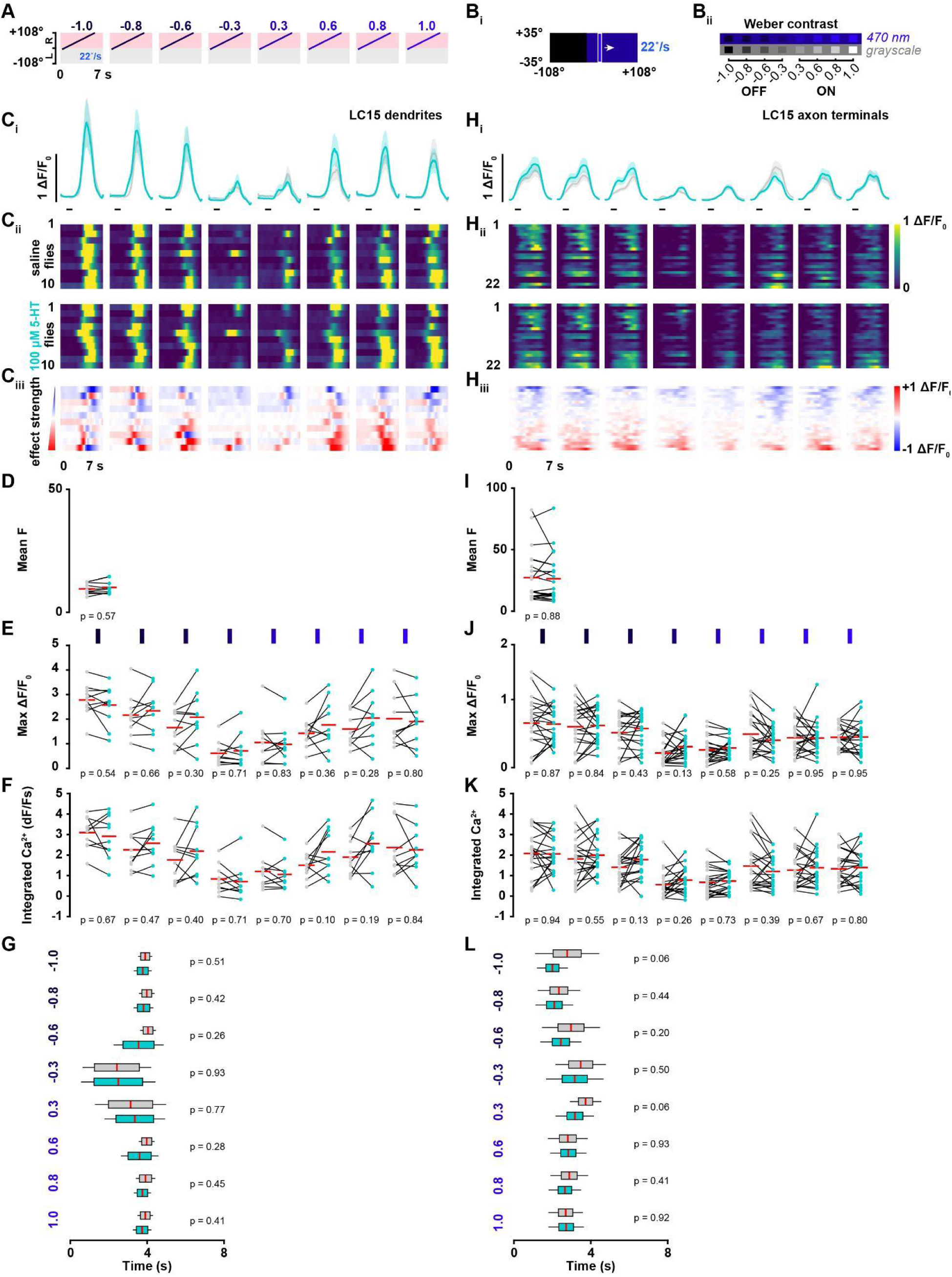
Dendritic and axonal LC15 responses show no effect of bath-applied 5-HT on luminance polarity. (A) Motion of the visual feature over time. The bar of varying luminance levels moved at a constant velocity of 22 °/s across the right side of the LED display. The Weber contrast ratio is given above. The mean irradiance values (light-OFF to light-ON) are 0.0, 0.009, 0.018, 0.039, 0.074, 0.092, 0.101, 0.11 μW m^-2^. (Bi) LED display schematic illustrating the 8.8° x 70° vertical bar stimulus presentation. (Bii) Weber contrast profile of the eight tested luminance levels. (Ci) Population mean GCaMP6f signal from the LC15 dendrites (± SEM shading) before (gray) and after 100 µM 5-HT (cyan). The horizontal scale bar (black) represents 1 second. (Cii) ΔF/F_0_ response raster plots displaying mean animal responses before (top) and after 100 µM 5-HT (bottom) (N = 10). Flies 1-10 (rows) are sorted the same in the pre-5-HT (top) and post-5-HT (bottom) condition rasters. (Ciii) Mean change in effect size rasters of LC15 dendritic GCaMP6f responses for each animal in the saline and 5-HT experimental groups. Animals (row) are sorted by increasing effect size strength. (D) Pairwise comparison of mean fluorescent intensity in the LC15 dendrites before (gray) and after 100 µM 5-HT (cyan). Horizontal red lines correspond to the population means. Two-tailed paired t-test, N = 10. (E) Pairwise comparison of maximum ΔF/F_0_ responses in the LC15 dendrites before (gray) and after 100 µM 5-HT (cyan). Two-tailed paired t-test, N = 10. (F) Pairwise comparison of integrated Ca^2+^ responses in the LC15 dendrites before (gray) and after 100 µM 5-HT (cyan). Two-tailed paired t-test, N = 10. (G) Pairwise comparison of time to half-max (s) for LC15 dendrite responses before (gray) and after 100 µM 5-HT (cyan). Two-tailed paired t-test, N = 10. (Hi) Population mean GCaMP6f signal from the LC15 glomerulus (± SEM shading) before (gray) and after 100 µM 5-HT (cyan). The vertical scale bar (black) represents 1 ΔF/F_0_. The horizontal scale bar (black) represents 1 second. (Hii) ΔF/F_0_ response raster plots displaying mean animal responses before (top) and after 100 µM 5-HT (bottom) (N = 22). Flies 1-22 (rows) are sorted the same in the pre-5-HT (top) and post-5-HT (bottom) condition rasters. (Hiii) Mean change in effect size rasters of LC15 glomerular GCaMP6f responses for each animal in the saline to 5-HT experimental groups. Animals (row) are sorted by increasing effect size strength. (I) Pairwise comparison of mean fluorescent intensity in the LC15 glomerulus before (gray) and after 100 µM 5-HT (cyan). Horizontal red lines correspond to the population means. Two-tailed paired t-test, N = 22. (J) Pairwise comparison of maximum ΔF/F_0_ responses in the LC15 glomerulus before (gray) and after 100 µM 5-HT (cyan). Two-tailed paired t-test, N = 22. (K) Pairwise comparison of integrated Ca^2+^ responses in the LC15 glomerulus before (gray) and after 100 µM 5-HT (cyan). Two-tailed paired t-test, N = 22. (L) Pairwise comparison of time to half-max (s) for LC15 glomerulus responses before (gray) and after 100 µM 5-HT (cyan). Two-tailed paired t-test, N = 22.

### Two-photon image analysis

Recorded imaging data was exported from SlideBook as 16-bit tiff image stacks and analyzed offline using custom written Matlab scripts. Images were corrected for motion artifacts and ROI selection was performed using a custom toolbox (39) (https://github.com/bjhardcastle/SlidebookObj). The ROI selection process was performed for each experimental video and identical for all recordings. Manually drawn ROIs were selected in accord with the anatomical region that was recorded. For glomerular recordings, a single ROI mask encompassed the whole glomerulus. For the dendrite recordings, the Ca^2+^ activity appeared as small fluorescent dots (lobula neurons perpendicular to the imaging plane) or small lines (parallel to the imaging plane). A toolbox function constructs an activity frame image (mean fluorescent intensity per pixel recorded during stimulus trials, subtracted by the mean of all extra-trial frames). All dendrites in frame that had a positive correlation between activity and the stimulus presentation were kept as individual ROIs. For every pixel in this ROI mask, the mean and standard deviation were calculated for the full time series.

### Fluorescence quantification

A time series was generated by calculating the mean fluorescence intensity of pixels within the ROI in each frame (F_t_). These mean values were then normalized to a baseline value as

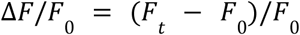

where F_0_ was the mean of F_t_ during the 0.5 s preceding stimulus onset. To compute the average time series across preparations with small variations in TTL synchronization, traces were resampled using linear interpolation. This procedure did not cause any detectable change in the original data.

The cumulative integral of each ΔF/F_0_ trace was obtained using the *cumtrapz* function in Matlab. The discrete integration of the GCaMP fluorescent intensity change over time is defined as

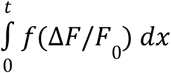

where the integrated Ca^2+^ measurement is the total area under the curve measured at the end of the stimulus period.

### FlyWire connectome analysis

We exported synaptic connectivity and neural morphology from the FlyWire connectome, using the FAFB v783 dataset (37). The EM reconstruction in v783 uses a coordinate system which is reversed from the fly’s anatomy, therefore the right optic lobe is labelled as the left hemisphere throughout the dataset. We used the following queries to generate data for Figure 7, and Tables 1-6.

- cell_type == LC15 && side == left
- cell_type == LC12 && side == left
- cell_type == 5-HTPLP01 && side == left
- nt_type == SER && output_neuropil == PVLP_L

Individual neuron IDs as well as the individual cells within the groups above were used in pathway analysis to assemble Figure 7. Cell type groups were analyzed for neurotransmitter profile, input and output neuropils, synaptic weight, and major upstream and downstream partners, presented in Tables 1-6.

## Results

We generated a set of visual stimuli to elicit responses in the two feature detectors. To assess figure-ground discrimination, a hallmark of some LCs, we used a ‘motion-defined’ bar composed of a narrow vertical bar that is randomly textured similar to the wide-field (WF) background, and thereby indistinguishable by contrast or luminance but rather only when it moves relative to the background (41). The motion-defined bar stimulus is highly engaging, stimulating saccadic pursuit in flight (42), and strongly activates high frequency sensitive T2a and T3 cells upstream of the LCs (43), but activates low frequency directional motion detectors T4 (ON edge-motion) and T5 (OFF edge-motion) less strongly (44).

We tested vertical motion-defined bars on stationary or moving backgrounds (Figure 1). We also tested a solid ‘luminance-defined’ bright (light-ON) or dark (light-OFF) bar on a uniform midlevel background. We tested bars of various luminance levels moving from the midline to the periphery at a constant speed (Figure 3). We tested OFF bars translating at different velocities (Figure 4) or cardinal motion directions (data not shown). We investigated the effects of bath-applied 5-HT on LC15, which increased the LC15 response to motion-defined bars translating across a stationary random texture background.

**Figure 4.**
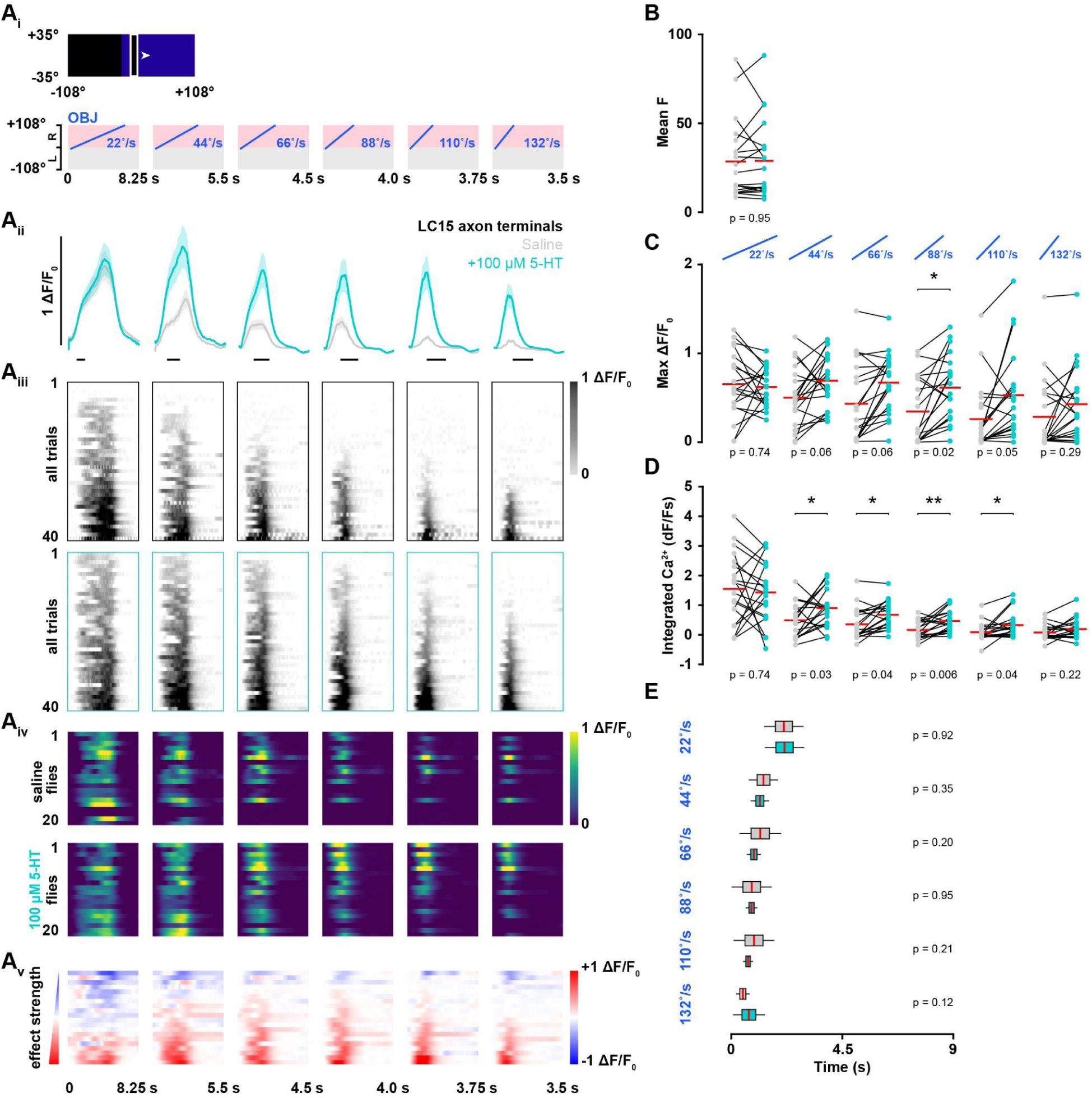
LC15 glomerular responses to OFF bar velocity are potentiated by 5-HT. (Ai) Schematic illustrating the 8.8° x 70° vertical bar presented on the LED display (top) and the motion of the translating feature over time (bottom). The OFF bar moved at six different velocities (22, 44, 66, 88, 110, and 132 °/s) across the right side of the LED display. (Aii) Population mean GCaMP6f signal from the LC15 glomerulus (± SEM shading) before (gray) and after 100 µM 5-HT (cyan). The horizontal scale bar (black) represents 1 second. (Aiii) ΔF/F_0_ response raster plots displaying individual trial responses before and after 100 µM 5-HT (cyan). Two repeats of each stimulus were shown to each animal (N = 20, n = 40). The trial responses are sorted by increasing peak ΔF/F_0_ response. (Aiv) ΔF/F_0_ response raster plots displaying mean animal responses before (top) and after 100 µM 5-HT (bottom) (N = 20). Flies 1-20 (rows) are sorted the same in the pre-5-HT (top) and post-5-HT (bottom) condition rasters. (Av) Mean change in effect size rasters of LC15 glomerular GCaMP6f responses for each animal in the saline to 5-HT experimental group. Animals (row) are sorted by increasing effect size strength. (B) Pairwise comparison of mean fluorescent intensity in the LC15 glomerulus before (gray) and after 100 µM 5-HT (cyan). Two-tailed paired t-test, N = 20. (C) Pairwise comparison of maximum ΔF/F_0_ responses in the LC15 glomerulus before (gray) and after 100 µM 5-HT (cyan). Two-tailed paired t-test, N = 20. (D) Pairwise comparison of integrated Ca^2+^ responses in the LC15 glomerulus before (gray) and after 100 µM 5-HT (cyan). Two-tailed paired t-test, N = 20. (E) Pairwise comparison of time to half-max (s) for LC15 glomerulus responses before (gray) and after 100 µM 5-HT (cyan). Two-tailed paired t-test, N = 20.

We performed calcium imaging in both the LC15 dendritic arbors and OG in the central brain (Figure 1Ai pink boxes, and inset). We targeted LC15 using the split-gal4 line OL0042B to drive expression of the calcium indicator GCaMP6f (Figure 1Aii) (27,38). Calcium imaging was performed in the right hemisphere of head-fixed flies (Figure 1Bi). We actively perfused temperature-controlled *Drosophila* saline throughout the duration of the experiment before switching the perfusate to 100 µM of 5-HT hydrochloride dissolved in *Drosophila* saline. Only flies that showed consistent responses to repeated stimuli in the control condition progressed to the 5-HT experimental condition and were included for analysis. Visual stimuli were presented on a curved LED panel display that covers 216° in azimuth (Figure 1Bii) (40). The anatomy of the LCs are shown in relation to the 5-HTPLP neuron (cyan) that provides 5-HT to the PVLP on the right hemisphere. The 5-HTPMP neurons (black) are also shown but do not innervate the PVLP (Figure 1Biii).

### 5-HT potentiates object motion responses by LC15

We displayed the same visual scene with four “figure-ground” stimulus conditions, randomly shuffled (Figure 1Ci). The first condition presents the WF moving rightward at 22°/s. The second condition presents the motion-defined vertical bar moving at 22°/s over the stationary WF. The third condition is a motion-defined bar held stationary +55° on azimuth while the WF background moves rightward at 22°/s. The fourth condition presents the bar moving opposite the background, both at 22°/s.

Figure 1Ci depicts the visual patterns as presented on the LED display (top) and the trajectory of the visual stimuli (bottom). Bath application of 5-HT was sufficient to cause a significant increase in the LC15 glomerular responses to a motion-defined bar over a stationary broadband background (Figure 1Cii). However, if the background was moving counter-directional beneath the bar, there was neither a response nor influence of 5-HT. Similarly, LC15 did not respond to a stationary motion-defined bar (+55°) defined by the rotation of the WF background behind it (Figure 1Cii), nor the broadband WF stimulus alone, and 5-HT had no effect (Figure 1Cii). A robust change in the glomerular responses is observable within the single trial responses after 5-HT bath application (Figure 1Ciii). We averaged the trial responses for each fly (Figure 1Civ), and observed a large degree of inter- and intra-animal variability. We lastly examined the effect size (post-5-HT subtracted from pre-5-HT) change for each animal between conditions, sorted by the magnitude of the change (Figure 1Cv). The effect size raster shows that the potentiation is confined to the stimulus that elicited visual responses from LC15, and the strength of the 5-HT effect is animal-specific.

We next tested whether the 5-HT application caused a tonic baseline potentiation of LC15 Ca^2+^ activity, which would be manifested by an increase in the mean fluorescence, F, but did not observe a significant change (saline = 26.0 ±21.0F; 5-HT = 25.5 ±20.6F) (Figure 1D). It was only during the acute presentation of the bar stimulus that we observed significant changes in both the maximum ΔF/F_0_ response (Figure 1E) and the time-integrated Ca^2+^ response (Figure 1F). We did not observe a significant change in the onset delay of visual responses, measured by time to half-max (Figure 1G), although there was considerable variability between preparations. These results indicate that 5-HT acts specifically upon the object detection gain of LC15, rather than on baseline excitation.

### 5-HT acts selectively on the presynaptic cell compartment - activating terminals not dendrites

To localize the site of gain modulation by 5-HT, we imaged visual responses in the postsynaptic dendrites in the lobula, reasoning that a change in gain of columnar inputs would modify the amplitude of GCaMP responses in the dendrites. Absence of 5-HT modulation would highlight compartment-specific feature computations and implicate central brain mechanisms acting within the axonal optic glomerulus. We first imaged dendritic responses in the lobula to the motion-defined bar stimuli (Figure 2Ai). LC15 cells are retinotopically arranged in the lobula and ROIs were drawn around the same posterior region of dendrite branches across preparations (see Methods). Although the visual responses were qualitatively similar in both cell compartments, i.e. largest response was to bar motion across the stationary ground (Figure 2A), we found no 5-HT potentiation in the LC15 dendrites (Figure 2Aii). Some individual dendrites in some individual trials displayed strong Ca^2+^ activity in response to the movement of the WF background (Figure 2A, columns one, three, and four). This is most likely due to phase locking of small-field responses to the spatial variation contained in the stimulus pattern, which is spatially averaged in the glomerular ΔF/F_0_ traces (Figure 1Cii). Note that visually activated GCaMP signals are much larger in a small group of dendrites than in the full population of axon terminals, which is also due to spatial averaging of time-shifted retinotopic terminals in the glomerulus. Unlike in the glomerulus (Figure 1), 5-HT did not modulate any of the visual response parameters we measured in the dendrites (Figure 2A-E), with the exception that some additional dendrites were recruited thereby increasing the number of ROIs collected in the post-5-HT condition (n_pre_ = 133, n_post_ = 168). But none of these measures were significant. These results suggest that 5-HT potentiation of LC15 responses is localized to the subcellular domain of the axon terminals within the glomerulus, where the individual LC15 clones converge and can be innervated by other circuits before conveying information to downstream pathways.

### 5-HT acts selectively on bar motion responses - does not modulate responses to varying object luminance

The directional motion vision system is split into ON and OFF subsystems (45), whereas feature detection pathways of the lobula so far studied are ON-OFF responsive (33,35), and LCs receive inputs from both systems (46–48). We therefore sought to test whether 5-HT selectively modulated ON or OFF selective pathways to LC15. We collected Ca^2+^ responses from the lobula dendrites (Figure 3C) as well as the central brain axon terminals (Figure 3H) while presenting an 8.8° x 70° vertical bar of eight different luminance values, set against a fixed luminance uniform background. The bar moved at 22°/s across the right side of the panel display (Figure 3A), with varying Weber contrast in randomized order (Figure 3Bi,Bii).

LC15 dendrites are contrast sensitive, but do not significantly prefer either polarity (Figure 3Ci), and these responses show no significant modulation by 5-HT in either individual or populations measures of effect strength, mean baseline fluorescence, max GCaMP amplitude, time-integrated GCaMP response, or onset delay (Figure 3C-G). The tonic level of GCaMP fluorescence (mean F) was higher in the glomerulus (control: 27 ±21F; 5-HT: 26 ±19F) than in the dendrites (control: 9.5 ±2.2F; 5-HT: 10.1 ±2.6F) but this is to be expected from averaging across all members of the cell class in volumetric imaging. Unlike the figure-ground stimulus (Figure 1), 5-HT did not modulate responses to object contrast in the axon terminals of the optic glomerulus (Figure 3H). Neither were any of the measured parameters significantly modulated (Figure 3I-L).

It is worth noting that mean peak ΔF/F_0_ values are larger for the solid bar than for the motion defined bar (compare Figure 2Aii second column with Figure 3Ci first column). This indicates that the dendritic GCaMP responses to the motion-defined bar in Figure 2 are not saturated or ‘clipped’. There is physiological ‘headroom’ in LC15 dendrites for 5-HT potentiation to have been manifested, but it is not. Taken together with the absence of contrast modulation suggests to us that 5-HT is unlikely to be acting upon ON or OFF presynaptic inputs to LC15 dendrites, but rather on the presynaptic terminals of LC15 itself in the glomerulus.

### 5-HT enhances speed sensitivity by LC15

The strong dendritic response to a dark bar (Figure 3C) motivated us to use this low frequency luminance-defined stimulus, rather than the high frequency motion-defined bar, to test whether 5-HT modulates LC15 responses to varying object velocity. We tested an 8.8° x 70° OFF bar moving at 22, 44, 66, 88, 110, 132°/s (Figure 4Ai) in randomized order. These bar velocities correspond to integer multiples of display pixels per second and were previously tested in the lab (35). Results from presynaptic glomerulus recordings are shown in Figure 4 and the corresponding results from postsynaptic dendrites can be found in Figure S1. LC15 responded best to the slowest-velocity OFF bar, and was attenuated with increased bar speed (Figure 4Aii). Speed attenuation was not observed in the dendrites (Figure S1), confirming previous findings (35). Remarkably, in the glomerulus, 5-HT delivery was accompanied by potentiated bar responses, especially at higher velocity (Figure 4Aii,Aiii) across animals in the experimental cohort (Figure 4Aiv,Av).

5-HT perfusion was not accompanied by a change in baseline GCaMP fluorescence, suggesting that enhanced velocity responses were not mediated by tonic Ca^2+^ accumulation (Figure 4B). The mean peak GCaMP response for the slowest 22°/s bar was unchanged, but the fastest velocities caused a statistically significant increase in peak ΔF/F_0_ at 88°/s (Figure 4C). The responses to the other high velocity bars were likewise increased however not individually significant (Figure 4C). The time-integrated Ca^2+^ responses showed that 5-HT significantly increased visually evoked Ca^2+^ accumulation for bars presented at 44, 66, 88, and 110°/s (Figure 4D). 5-HT did not significantly affect the onset delay of the visual response (Figure 4E). None of these significant changes were observed in the mean lobula dendrites (Figure S1B,C). As with figure-ground discrimination (Figure 1), 5-HT induced velocity gain is localized to the synaptic outputs implicating neuromodulatory targets within glomerular circuits of the CB rather than within retinotopic inputs from the OL.

### A similar bar sensitive feature detector, LC12, is not modulated by serotonin

We next sought to test whether 5-HT has similar effects on other LC types, and examined another bar motion sensitive detector, LC12 (Figure 5A). As with LC15, we did not observe any change in baseline fluorescence after 5-HT application in LC12; neither did we observe any modulation of visual responses. Figure 5B shows the results from the motion-defined bar over WF background experiment used in Figures 1 and 2. We observed low amplitude responses in the LC12 glomerulus to the motion-defined bar, but no potentiation from 5-HT application (Figure 5B). We measured a subtle yet statistically significant *reduction* in the mean ΔF/F_0_ response to the third stimulus condition, a static bar defined by the WF, after 5-HT bath application (Figure 5Ci). The integrated Ca^2+^ response for this stimulus sequence was significantly reduced as well (Figure 5Cii). We tested bars of varying luminance levels (Figure 5D), or velocities (Figure 5E), but did not observe any changes after 5-HT application (Figure S2). Taken together, our recordings from LC12 and LC15 indicate that serotonergic modulation of visual responses is cell type specific even within the family of structurally similar feature detectors and modulation acts selectively in the presynaptic compartment.

**Figure 5.**
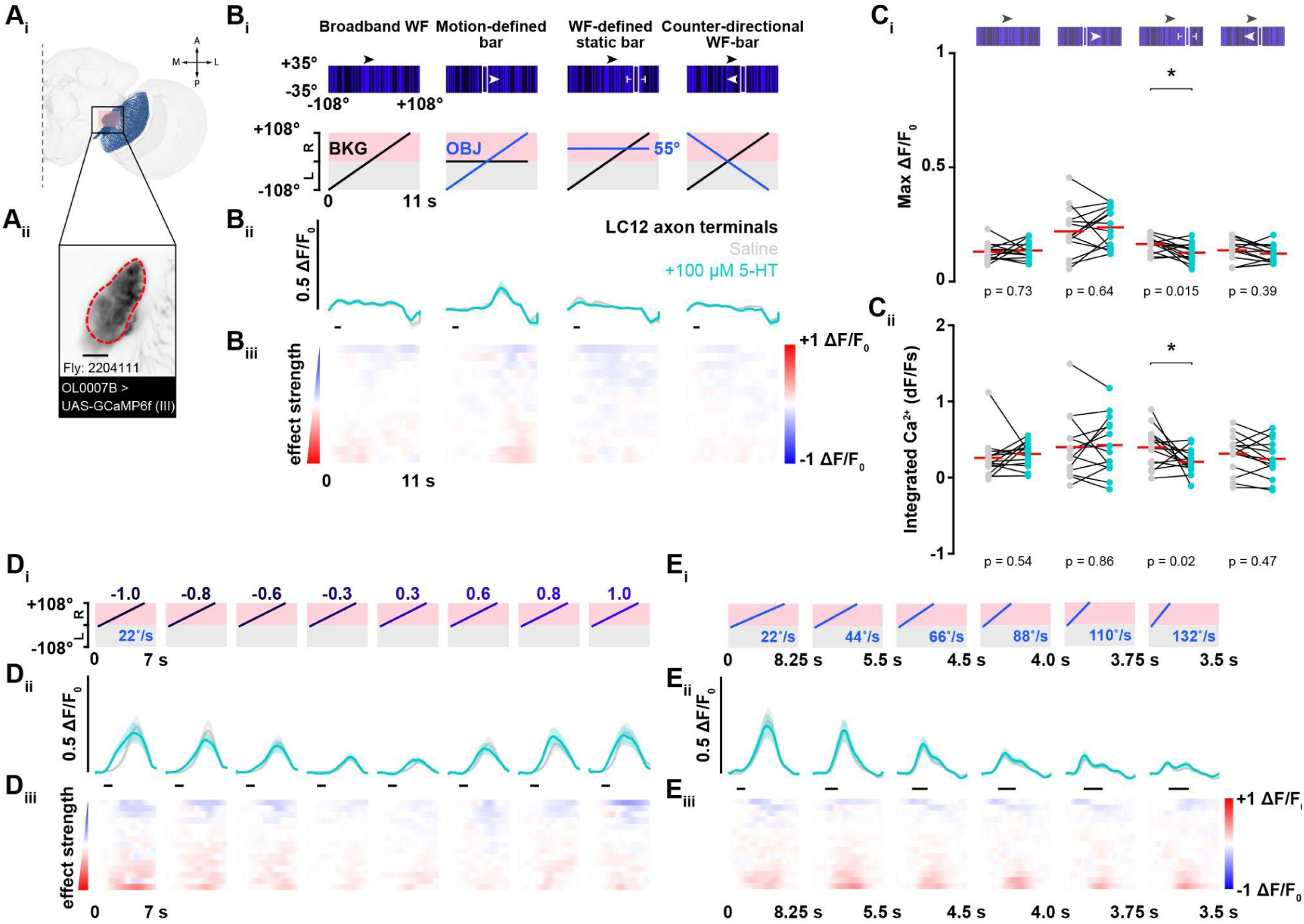
The LC12 glomerulus does not show potentiation to translating vertical bars after 5-HT bath perfusion. (Ai) FlyWire connectome reconstruction of LC12 cells (blue) in the right hemisphere. The pink box represents the imaging site of the LC12 glomerulus. The LC12 dendritic projections receive signals in the lobula and send that information downstream to the glomerulus in the VLP. (Aii) Two-photon fluorescent intensity image showing the glomerulus, or axonal bundle of LC12 cells. The red dotted line defines the region of interest. Scale bar, 25 µm. (Bi) Schematic showing the visual stimulus presented on the LED display (top) and the motion of the visual feature elements over time (bottom). The second column displays the LC12 glomerular response to a motion-defined equiluminant vertical bar over a stationary broadband background before (gray) and after bath perfusion of 100 µM 5-HT (cyan) in saline. (Bii) Population mean GCaMP6f signal from the LC12 glomerulus (± SEM shading) before (gray) and after 100 µM 5-HT (cyan). (Biii) Mean change in effect size rasters of LC12 glomerular GCaMP6f responses for each animal in the saline and 5-HT experimental groups. Animals (row) are sorted by increasing effect size strength. (Ci) Pairwise comparison of maximum ΔF/F_0_ responses in the LC12 glomerulus before (gray) and after 100 µM 5-HT (cyan). Two-tailed paired t-test, N = 14. (Cii) Pairwise comparison of integrated Ca^2+^ responses in the LC12 glomerulus before (gray) and after 100 µM 5-HT (cyan). Two-tailed paired t-test, N = 14. (Di) Motion of the visual feature over time. The bar of varying luminance levels moved at a constant velocity of 22 °/s across the right side of the LED display. (Dii) Population mean GCaMP6f signal from the LC12 glomerulus (±SEM shading) before (gray) and after 100 µM 5-HT (cyan). (Diii) Mean change in effect size rasters of LC12 glomerular GCaMP6f responses for each animal from the saline to 5-HT condition. (Ei) The motion of the translating feature over time. The OFF bar moved at six different velocities (22, 44, 66, 88, 110, and 132 °/s) across the right side of the LED display. (Eii) Population mean GCaMP6f signal from the LC12 glomerulus (±SEM shading) before (gray) and after 100 µM 5-HT (cyan). (Eiii) Mean change in effect size rasters of LC12 glomerular GCaMP6f responses for each animal in the saline and 5-HT experimental group.

### Serotonin modulates speed sensitivity in the high-velocity regime

After observing velocity-dependent changes to LC15 glomerular responses with 5-HT bath application (Figure 4), we calculated the visual response gain induced by 5-HT, i.e. the ratio of peak Ca^2+^ responses in the 5-HT condition relative to the saline control, for four experimental groups (Figure 6). The LC12 glomerulus does not show a significant 5-HT gain change at any of the tested bar velocities (Figure 6, blue) compared to the control saline treatment (gray). By contrast, LC15 axon terminals in the glomerulus show gain enhancement that peaks in the midrange of tested velocities (cyan). This gain enhancement is not observed in the lobular dendrites (Figure 6, purple), where response gain is no different from saline controls imaged in the glomerulus. These data indicate that LC15 gain modulation is tuned for intermediate object speeds, which could support social behaviors that occur within this range of angular velocities (49).

**Figure 6.**
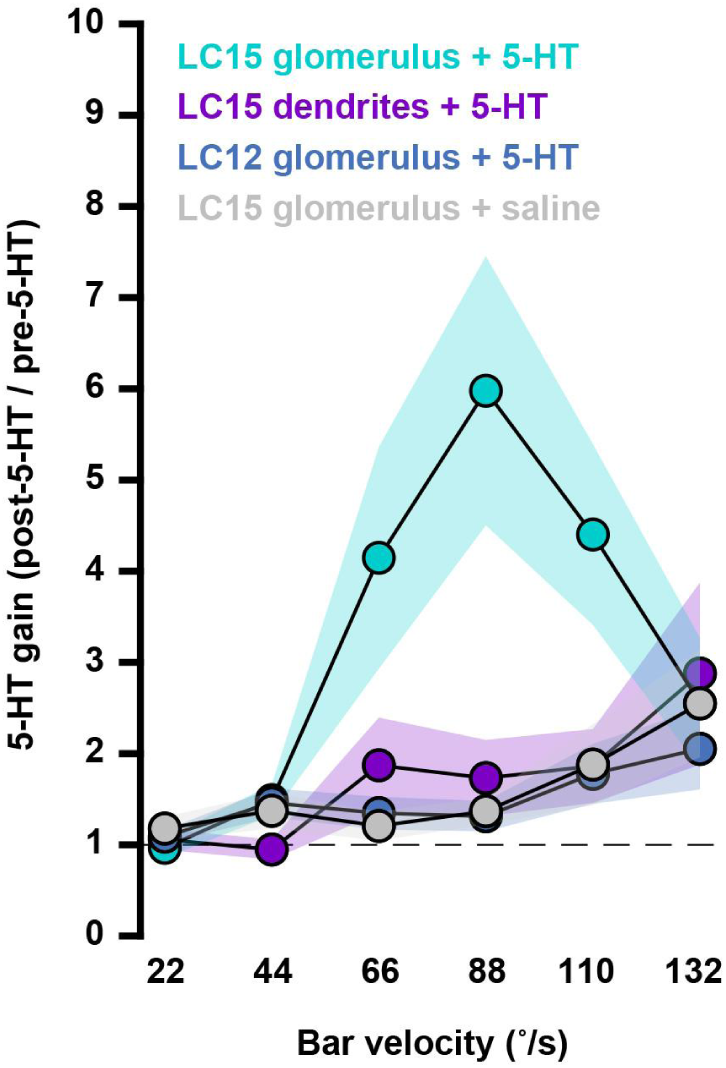
5-HT enhances speed tuning in LC15 glomerulus. The relationship between the mean ratio (+SEM) of individual max ΔF/F0 responses to six OFF bar velocities. Four post-/pre-groups are shown: LC15 glomerular responses after 100 μM 5-HT (cyan), LC15 dendritic responses after 5-HT (purple), LC12 glomerular responses after 100 μM 5-HT (blue), and LC15 glomerular responses after saline (gray). LC15 glomerulus + 5-HT, N = 20; LC15 dendrites + 5-HT, N = 10; LC12 glomerulus + 5-HT, N = 14; LC15 glomerulus + saline, N = 10. The dashed horizontal line denotes no change.

### Connectomics reveal multiple inhibitory circuit motifs with putative serotonergic LC15 interactions

We used the FlyWire EM reconstruction connectome database to examine the chemical synaptic partners of LC15 (37). This resource labels all of the synaptic partners in the brain but, unfortunately, cannot identify extra-synaptic paracrine partners involved in volumetric hormonal neuromodulation. The connectome does however provide us with a wealth of information about the participating neurons including the neural architecture of the two feature-processing VPNs studied here. First, we examined the connectome for any shared upstream inputs to both LC12 and LC15, starting with the lamina monopolar neurons L1-L5. We also identified the downstream partners of these two LCs for potential serotonergic intersections.

The major inputs by synaptic count to LC15 include TmY4, Li27, TmY9q⊥, TmY9q, Li31, PVLP101b, CB0346, Tm25, PVLP103, and LmA2. TmY4, TmY9q⊥, and TmY9q are three horizontally-oriented neurons predicted to prefer OFF objects along their axis of orientation. Other strong LC15 disynaptic pathways include T3, a small-object detector, and other object-detecting cells, Tm25 and Tm21 (50). This report also showed that three populations of glutamatergic neurons (Dm3q, Dm3p, Dm3v) play a critical role in filtering OFF activated signals supplied to LC15. TmY4, TmY9q⊥, TmY9q, Li30, and Tm25 are cholinergic. PVLP103, CB0346, PVLP101b and CB0140 are GABAergic. Li27 is glutamatergic (Table 1). The major outputs from LC15 by synaptic count are PVLP121, CB0381, CB2049, PLP115_b, CB0346, PVLP028, PVLP118, LT61b, PVLP151, and PVLP018 (Table 2).

The major inputs to LC12 are Tm3, Tm4, PVLP097, T2, TmY3, Li17, TmY11, cL21, T3, and Tm2. The connectome shows that L2 synapses onto Dm17 which connects to TmY3. L2 also synapses onto Tm4 (implicated in the OFF motion pathway), which has connections to Li31, a GABAergic inhibitory neuron that crosses the midline of the brain and connects to the LC15 glomerulus on the contralateral hemisphere. Tm3, Tm4, T2, TmY3, TmY11, T3, and Tm2 are cholinergic. PVLP097, Li17, and cL21 are GABAergic (Table 3). The major outputs from LC12 by synaptic count are PVLP097, PVLP025, PVLP013, PVLP037, CB0813, CB1340, cL21, PVLP120, PVLP100, and AVLP152 (Table 4). These results indicate that LC12 and LC15 share no major zero hop presynaptic inputs, and only a single one-hop input from L5. L5 itself may express 5-HT7 but this has yet to be confirmed (21).

Figure 7A displays a connectivity diagram between the top ten pre- and postsynaptic partners to LC15 and LC12, stratified by neuropil from the optic lobe into the central brain. The top ten pre- and postsynaptic partners to LC15 and LC12 are also shown and colored by their putative neurotransmitter type. It should be noted that this is a non-exhaustive list meant to highlight similarities or differences in the chemical synaptic wiring between these two bar-sensitive object detectors. We also queried the database for synapses between serotonergic 5-HTPLP01 and our LCs of interest or their immediate partners, but we did not identify any. Interestingly, 5-HTPLP01 may co-release 5-HT volumetrically and glutamate synaptically (see (52–54)). 5-HTPLP01 does connect indirectly to LC15 through a centrifugal GABAergic interneuron, cL21, which provides feedback inhibition to LC12 and synapses onto another set of GABAergic neurons, LmA2. LmA2 connects to LC15 dendrites at the lobula (Figure 7A, Figure S4B). Tables 5 and 6 list the top ten pre- and postsynaptic partners for 5-HTPLP01. The strongest synaptic input to 5-HTPLP01 comes from ALVP169, cholinergic cells that make reciprocal synapses onto 5-HTPLP01 from the AVLP, but 5-HTPLP01 also receives GABAergic input from CB0534 and AVLP081, and glutamatergic input from PVLP106 (Figure S3A,S3C).

**Figure 7.**
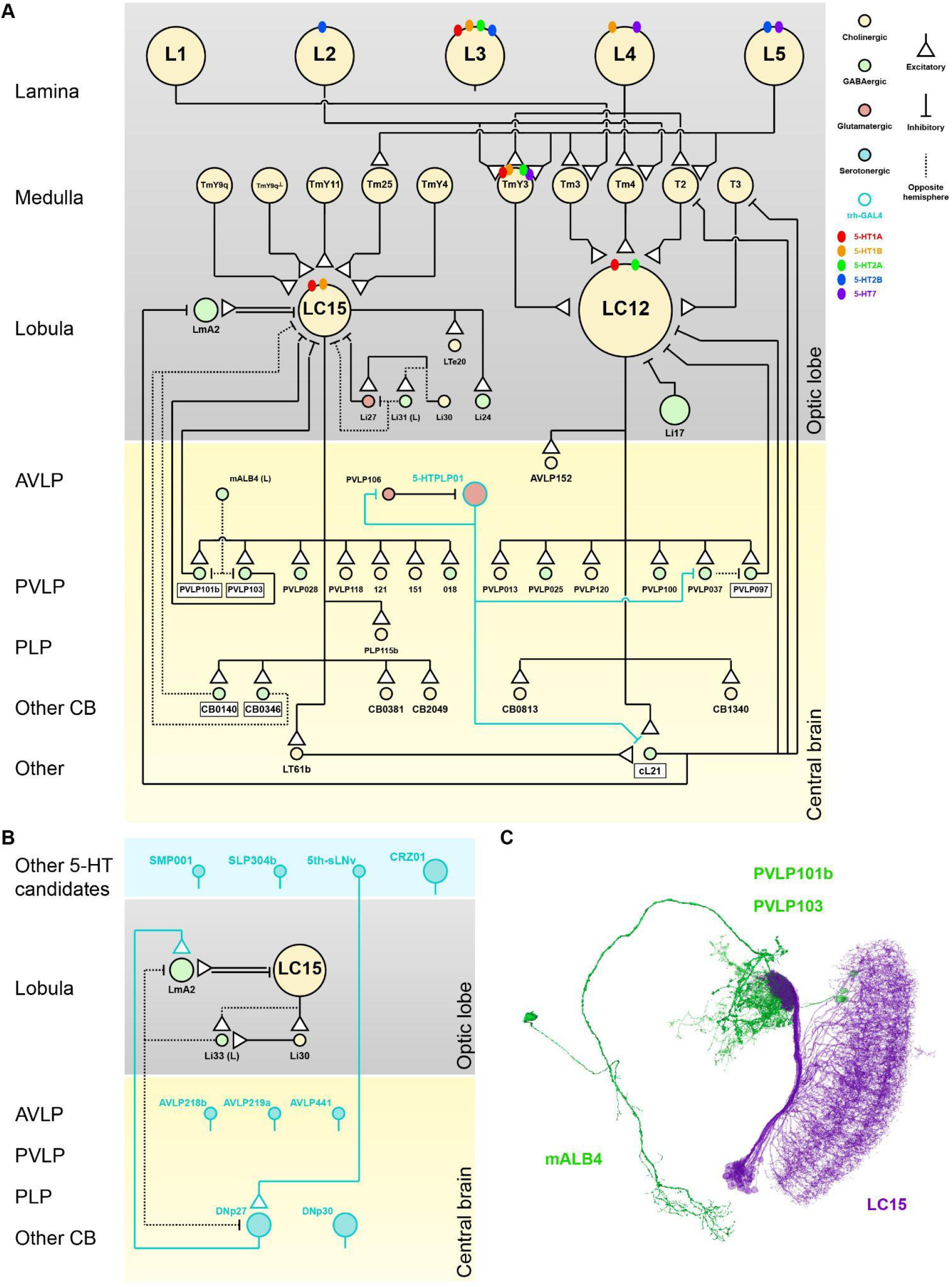
Connectomics highlights specific circuit motifs and candidate sites for 5-HT neuromodulation. (A) Wiring diagram of LC15 and LC12 illustrating the connections between the major presynaptic and postsynaptic partners and the synaptic connections to 5-HTPLP (37). Cells are colored by neurotransmitter type and are stratified by neuropil. The colored dots detail the 5-HTR expression where available. Bordered cell names refer to neurons that both send and receive synapses from LC15 or LC12. The 5HTPLP01 is labelled by trh-GAL4 (cyan) but uses glutamate as its neurotransmitter so the connections to downstream partners are shown as inhibitory. 5-HTPLP01 makes reciprocal connections to another glutamatergic neuron, PVLP106. 5-HTPLP01 makes the largest number of synapses onto cL21 (703, 6%) but otherwise does not make connections to the members of the LC15 pathway. cL21 is a pair of centrifugal GABAergic cells that have input synapses in the PVLP and outputs in the lobula. Both PVLP106 and cL21 receive most of their input synapses from LC17 (3,743, 54% and 4,560, 25%). 5-HTPLP01 does synapse onto a downstream target of LC12, a GABAergic cell class called PVLP037. PVLP037 has 4 cells per hemisphere and makes 684 inhibitory synapses onto PVLP097, a recursive partner of LC12. 5-HTPLP01 sends and receives most of its connections from AVLP interneurons. (B) The neurons of the right hemisphere that have output synapses in the PVLP and use 5-HT as a neurotransmitter (cyan). Direct connections to the LC15 VPN network are also shown. The only serotonergic cell that synapses onto members of the LC15 pathway is DNp27. DNp27 is a large bilateral descending neuron that innervates the right SMP, PLP, PVLP and AVLP. Output synapses are observed in the central brain and in the lobula but presumably most outputs are located in the VNC (tectulum, lower tectulum, and abdominal segment, see (51)). In the lobula, DNp27 makes synapses onto a population of 17 lobula-medulla amacrine cells, LmA2. The LmA2 cells make up a patchwork tiling the visual field and the LmA2 cells make 583 synapses onto LC15 in the lobula. (C) Morphology of the mALB4-PVLP101b-PVLP103-LC15 circuit element. LC15 (purple) and the GABAergic partners (green) that interact at the LC15 glomerulus. mALB4 is a GABAergic neuron that has input synapses in the PVLP and the SEZ. The output synapses are solely located in the PVLP, where it connects to PVLP101b and PVLP103. PVLP101b contains two GABAergic cells that have reciprocal (input and output) synapses with LC15. PVLP103 similarly contains two GABAergic cells that receive 341 synapses from LC15 and makes 593 synapses back onto LC15.

#### I. Peripheral optic lobe

We queried the FlyWire connectome database to identify any serotonergic neurons (i.e. neurons that use 5-HT as a neurotransmitter) that interact with LC15. We observed that only one, the expansive serotonergic descending neuron, DNp27, makes synaptic connections onto GABAergic LmA2 neurons in the lobula, which in turn synapse onto LC15 (Figure 7B) (see (51)). DNp27 receives inputs from another serotonergic neuron, 5th-s-LNv which is a member of the evening oscillator in the circadian clock circuitry and required for a bimodal sleep-activity rhythm (55–58). Figure 7B displays the connectivity between all serotonergic neurons with synaptic output to the numerically top ten pre- and postsynaptic partners to LC15 and LC12, stratified by neuropil from the optic lobe into the central brain. When the 5th-s-LNv is activated, this could enhance responses from DNp27, to increase the GABAergic tone in the lobula through the LmA2 amacrine cells at a particular time in the circadian rhythm.

This analysis uncovered another lobula circuit element where lateral inhibitory neurons in the lobula interact directly, or indirectly through the glutamatergic lateral neurons, Li27. There are two Li27 neurons per hemisphere that form numerous synapses onto the dendrites of LC15 (1,670 synapses to all LC15 cells, 24% of all inputs). Li27 receives input from TmY4, TmY9q⊥, and TmY9q as well as a GABAergic interneuron, Li31. Li31 receives cholinergic inputs from Tm4 and Tm3 but also from a lateral neuron, Li30 (Figure S4A) before synapsing onto Li27 and LC15. Li30 is a cholinergic cell upstream of Li31 which receives most of its synapses from Y3. These three cells are positioned to act as a gain control mechanism upon LC15 in the lobula, independent of the two glomerular motifs downstream. The glutamatergic Li27 cells may provide shunting inhibition of LC15 to filter out noise from object stimuli.

LC15, unlike LC12, shows a large portion of output synapses in the lobula (∼10%) (35,37). 818 of the 2945 synapses from LC15 cells connect to LT61b (28%, ACh), 379 synapses to LTe20 (13%, ACh), 639 to LmA2 (22%, GABA), and 326 to Li24 (11%, GABA). These four partners receive ∼75% of the total number output synapses in the lobula from LC15. These following cells roughly make up the remaining 25%. 3%: LC21, LC31a; 2%: LC9, LC31c, Tm31; 1%: LTe12, Li05, Li06, LmA5, LC11, LC28a, LC31b, LPLC1, Tm5f, and Tm35. The partner with the highest synaptic weight, LT61b, is a single cell per hemisphere (Figure 7A; Figure S4B) and it only innervates the lobula at layer 2. LmA2 is a patchwork of 17 GABAergic cells that have synapses in both layer 2 and layer 4. LTe20 also has 1 cell per hemisphere and only makes synaptic connections with LC15 in layer 2. Lastly, the 4 intrinsic Li24 cells per hemisphere receive synapses from LC15 in both layers 2 and 4. This demonstrates that the GABAergic cell groups (LmA2 and Li24) receive synapses from LC15 in lobula layers 2 and 4 while the cholinergic partners are single-cells (LT61b and LTe20) that synapse only in layer 2 and bypass the LC15 glomerulus.

#### II. Central brain

Two glomerulus-specific circuit motifs were also uncovered: 1) contralateral suppression and 2) gain control through release from inhibition. The wiring diagram suggests that the neural architecture exists to provide contralateral GABAergic suppression in LC15. Downstream partners of LC15, CB0140 and CB0346 are GABAergic cells that innervate the LC15 glomerulus and synapse onto the other LC15 glomerulus (Figure S5B). This axo-axonal suppression appears similar to what has previously been reported in another different LC, LC6 (59).

Our connectivity analysis also shows that LC15 makes connections to PVLP101b and PVLP103 in the OG. These GABAergic lateral inhibitory neurons also make reciprocal synapses back onto LC15 and presumably provide the majority of the inhibitory tone at the LC15 glomerulus through feedback inhibition. PVLP101b and PVLP103 both share an upstream partner, mALB4, a GABAergic neuron from the left hemisphere that synapses at the LC15 glomerulus but does not make direct connections to the LC15 cells themselves. The dendritic arbor of the mALB4 neuron also innervates the subesophageal zone, SEZ (Figure 7C) (Figure S5A). LC12 makes reciprocal synapses with a population of lateral inhibitory neurons, PVLP097. There are 7 GABAergic PVLP097 cells per hemisphere that make reciprocal synapses with LC12. PVLP097 is LC12’s highest output partner (15,920, 28%), and LC12 is PVLP097’s highest output partner (2,403, 25%). PVLP037 constitutes 4 GABAergic neurons upstream of PVLP097 that cross hemispheres (794, 4%). 5-HTPLP makes glutamatergic synapses onto PVLP037 (98, 1%) (Figure S6).

## Discussion

### Receptor expression does not predict neuromodulation of visual responses

As in vertebrates, multiple serotonin receptor (5-HTR) subtypes have been identified in *Drosophila* that initiate both excitatory and inhibitory GPCR-mediated functions (60–63). Five subtype receptors coupled to identified GPCRs have been identified in *Drosophila* (5-HT1A, 5-HT1B, 5-HT2A, 5-HT2B, and 5-HT7) that cause specific metabolic cascades through second messenger systems. Two of these receptors inhibit cAMP synthesis (5-HT1A, 5-HT1B), decreasing intracellular Ca^2+^ levels. The other three either promote cAMP production through activation of adenylate cyclases (5-HT7), or mediate hydrolysis of inositol phosphates (5-HT2A, 5-HT2B) leading to increased intracellular Ca^2+^ levels (64,65). 5-HTR expression was distinct for each receptor subtype in the adult brain neuropils including the mushroom bodies (MB), central complex (CX), optic lobes (OL) and antennal lobes (AL) (7). The 5-HTR expression profile is different between LC12 and LC15; LC15 displays high levels of 5-HT1A and 5-HT1B (66); YZ Kurmangaliyev, *personal communication*). Both 5-HT1A and 5-HT1B are coupled to the G_i/o_ GPCR which reduces cytosolic cAMP through the inhibition of adenylate cyclase (AC) activity. Immunohistochemistry analysis revealed that LC12 shows high levels of 5-HT1A and 5-HT2A in the presynaptic axon terminals of the glomerulus but not the postsynaptic dendrites in the lobula. 5-HT2A receptors are coupled to the GPCR G_q/11_, and activation leads to the hydrolysis of inositol phosphates and a subsequent intracellular Ca^2+^ increase. Sparse synaptic labeling of 5-HT suggests that the LC glomeruli in the VLP receive most of their serotonergic modulation via volumetric release of the neuromodulator (66).

A class of columnar neurons upstream of the LCs, lamina monopolar L2, show 5-HT sensitivity. L2 cells express 5-HT1A and 5-HT2B, and upon administration of exogenous 5-HT, showed a gradual increase in basal Ca^2+^ fluorescence in medulla layer M2 over the 6-minute time frame. This effect was dramatically reduced in 5-HT2B^-/-^ mutant flies. 5-HT2B, like 5-HT2A, are coupled to G_q/11_, suggesting that 5-HT activation of 5-HT2B on L2 neurons may enhance the response to OFF objects (i.e. objects darker than the background). This report suggests that the 5-HT2B receptor is necessary for serotonergic potentiation of L2 Ca^2+^ responses observed in the lamina (21). L2 is an upstream input for LC12 but not LC15 and we show that these two VPN populations do not share any common partners directly upstream (Figure 7A). It is possible that a feed-forward modulatory effect of 5-HT on L2 potentiates OFF responses in downstream partners, like LC12. Lastly, neither LC12 or LC15 express 5-HT7, but recent work has shown that 5-HT7-expressing GABAergic LNs in the antennal lobe mediate subtractive gain control in the PNs of the CO_2_-sensitive V glomerulus (7). In these inhibitory cells, 5-HT7 activates the Gs protein, which stimulates AC, increasing intracellular cAMP (15,67,68).

Here, we show that 5-HT modulation is highly selective within the LC feature detection pathway. The effects of 5-HT application are (i) stimulus-specific, enhancing responses to bar motion but not ground motion (Figure 1); (ii) they are anatomically compartmentalized, appearing in the axon terminals but not the dendrites (Figure 2); (iii) selective for visual stimulus parameters, as 5-HT application has no influence over luminance tuning (Figure 3), yet boosts response amplitude at different stimulus velocities (Figure 4); finally, (iv) the effects of 5-HT application are cell-type specific, modulating LC15 but not LC12 (Figure 5).

These results are unexpected given that LC15 purportedly expresses 5-HTRs that inhibit adenylate cyclase (64); YZ Kurmangaliyev, *personal communication*). By contrast, we expected to observe a potentiation effect in LC12. What is the role of excitatory 5-HT2A receptors on LC12 cells if not for modulating visual responses or to increase basal Ca^2+^? 5-HT is involved in a multitude of physiological mechanisms and behavioral phenotypes. 5-HTR expression has been shown to be involved in ‘higher order’ biological functions such as the entrainment of circadian rhythm, aggression, and courtship circuitry, or cell metabolic functions where 5-HT may play a role (8,9,57).

### Serotonin bath application, and presumably volumetric release in the VLP, reveals a coordinated increase in the feature-detecting parameter space

Bath application of any neuromodulator is experimentally convenient and is the only way to demonstrate the specificity of the signal rather than indirect action via another neuromodulator. Under normal conditions, endogenous 5-HT likely fills the ventrolateral protocerebrum (VLP) *via* volumetric release through 5-HTPLP01 to cause non-cell-autonomous neuromodulatory effects on lobula columnar (LCs) cells (1,66,69). This serotonergic activity in the VLP is likely to be extra-synaptic since 5-HTPLP is glutamatergic (Figure 7A). Our results are therefore notable in that the effects of bath application come on quickly, are object-stimulus-specific, cell-type-specific, and subcellular domain-specific (Figure 1-5). Indeed, there was no stimulus parameter that was shared across any response dimension we could find. Our results also suggest that neuromodulatory changes in VPN responses are not simply controlled through activation of volumetric neurons but through iLN activation that leads to specific downstream release from feedback inhibition. Previous behavioral experiments performing optogenetic targeting of trh-gal4 showed that a rigidly-tethered fruit fly tracks a bar significantly better during LED stimulation but had no effect on the small-object response (70). The trh-gal4 promoter line genetically targets cells expressing tryptophan hydroxylase (Trh) which is the rate-limiting enzymatic step in the production of 5-HT (71). These activation experiments (trh-gal4 > UAS-CsChrimson) are unlikely to fully capture the effect of state-dependent changes in sensorimotor transduction unless the inhibitory partners are also controlled. There is no way of knowing from these results if 5-HT is specifically causing this phenotype; it could result from downstream partners of serotonergic neurons instead of non-synaptic hormonal release (Figure 7B). To best mimic the neuromodulatory conditions for the LC15 glomerulus, optogenetic targeting of trh-gal4 serotonin-releasing neurons was insufficient to be used alone. The most parsimonious explanation is that 5-HT is acting extra-synaptically on a subset of glomerulus-specific inhibitory partners to enhance select visual responses in LC15 (Figure 7). The 5-HTR profiles of these lateral inputs are currently unknown.

### Identifying feedforward and feedback potentiation of feature detection

In the VLP, circuit-level potentiation may include increasing the gain of opposing cell classes (e.g. cholinergic and GABAergic cells) to allow for changes in the visual feature parameter space or SNR. Bath applications of 5-HT in brain explants result in increased Ca^2+^ activity in both GABAergic 5-HT1A receptor-expressing neurons, as well as cholinergic short neuropeptide F receptor and 5-HT1A receptor-expressing neurons (72). 5-HT may change the activity or responsiveness of these inhibitory cells, thereby affecting the GABAergic tone over LC15. If not through the direct effects of 5-HT onto the LC15 cells themselves, the potentiation of LC15 by 5-HT must be caused in a non-autonomous manner by a change in another element of the visual feature detection pathway. The two most likely non-autonomous ways for this potentiation to take place would be through 1) feedforward excitation, or 2) release from feedback inhibition. Feedforward excitation would involve serotonergic potentiation of excitatory inputs and/or decrease in inhibitory inputs upstream of LC15 in the optic lobe to increase the ‘gain’. Release from feedback inhibition would be characterized by top-down inhibition of GABA-releasing inhibitory cells intrinsic to the LC15 glomerulus.

If potentiation in LC15 were predominantly controlled by feedforward excitation, we reason that 5-HT would act upon presynaptic inputs to LC15 expressing excitatory 5-HTRs. Here, 5-HT would increase cholinergic activity upstream of LC15, presumably through TmY cells. Inhibitory 5-HTRs might also be located on Dm cells which synapse onto LC15 in the lobula. The potentiated responses would be relayed from the lobula to the glomerulus. We did not observe significant potentiation of LC15 responses in the lobula to any stimulus after 5-HT. Our dendrite recordings also support previous research showing a lack of velocity-tuned responses in the dendrites (26,35). We did identify two cholinergic partners that receive output synapses from LC15 in lobula layer 2, LT61b and LTe20. LT61b is a single-cell that has its output synapses in the ipsilateral lobula, the ipsilateral and contralateral AVLP and PVLP. This neuron partners with over 300 cells downstream including cL21 in the central brain. LTe20 is also a single neuron that has outputs in the lobula and PVLP and makes a total of ∼3,400 output synapses with >200 synapses onto ipsilateral LmA2 cells and ∼150 connections to Li24 neurons (Figure 7A).

If potentiation in LC15 were caused by release from feedback inhibition, this would require evidence of inhibitory cells that send and receive synapses to LC15. These partners must also be contained to the cellular subdomain of the OG, and that the OG are under some degree of active inhibitory control. 5-HT would disinhibit specific glomeruli in two ways (i) 5-HT inhibits GABAergic partners at the LC15 glomerulus (via 5-HT1A, or 5-HT1B), or (ii) 5-HT excites inhibitory GABAergic cells that themselves synapse onto inhibitory partners of LC15 (5-HT2A, 5-HT2B, or 5-HT7). Our experimental results suggest that in LC15, the individual retinotopically-arranged dendrites in the lobula are pooled together at the OG and that 5-HT has direct effects on the excitability of the presynaptic partners in the VLP. Through cotransmission, 5-HTPLP01 may inhibit downstream synaptic partners with glutamate signaling, and through hormonal release of 5-HT, activate others in the PVLP. 5-HT release potentiates responses in the LC15 glomerulus to select context-specific features while strengthening the filter through active inhibition. This increase in sensitivity (LmA2 release from inhibition via cL21, Figure 7B) is temporally linked with the release of 5-HT from 5-HTPLP01. Neuromodulators like 5-HT may disinhibit specific OGs, allowing responses to higher spatial and temporal frequencies, increasing response strength and sensitivity in the VLP, thereby extending the dynamic range of the feature detector. This phasic effect likely fluctuates with the internal and behavioral state of the animal, especially during changes in locomotion and escape behavior (69,73). Serotonin unexpectedly caused a selective Ca^2+^ increase in only the faster velocity OFF bars, tuning object movement velocity gain (Figure 4D, Figure 6). Angular velocity information about an expanding loom and its functional relevance to takeoff behavior has been shown in the looming-sensitive LC4 (74,75). Taken together with our findings suggest that presynaptic VPN responses can be sculpted and modulated at the OG, and the LC receptor expression profile is not necessarily indicative of that cell’s neuromodulatory potential.

### Similarities to olfactory feature detection

Our findings suggest functional similarity to the antennal lobe, another glomerular neuropil of the *Drosophila* brain that encodes olfactory features (65). The effects of 5-HT on the olfactory receptor neuron (ORN) to projection neuron (PN) synapse and on lateral GABAergic inhibition has been much closer examined here (18,19,76). The antennal lobe, the functional equivalent of the mammalian olfactory bulb, has a diversity of GABAergic local neurons (LNs) that interact with the ORN and PN cells, shaping activity (77–88). The similarities in anatomical architecture between these two neuropils have previously been noted (89,90).

The mAL neurons are *fru+* sexually-dimorphic GABAergic cells located in the medial antennal lobe that have been implicated in pheromone sensing and aggression (91,92). Activation of mAL significantly increases intermale aggression and the phenotype was significantly reduced when GABA release was attenuated in mAL (93). Different mAL neurons have been shown to be necessary for maintaining a ‘gain control’ mechanism over P1 neurons and activation of mAL neurons have been shown to suppress male courtship behavior (94). The mALB4 neuron, similarly to mAL, has input synapses in the GNG but instead of ramifying in the superior protocerebrum, mALB4 sends 344 inhibitory synapses onto partners in the PVLP where it could operate as a gain control mechanism across multiple OG in the PVLP (Figure 7C). The 5-HTR expression profile of mALB4 is unknown but, as a GABAergic mAL neuron, expression of 5-HT7 suggests that 5-HT would boost responses in these inhibitory neurons (7). If 5-HT is available in the PVLP extracellular space, mALB4 activation would strongly inhibit PVLP101b and PVLP103, thereby releasing LC15 from feedback inhibition specifically at the glomerulus (Figure 7A). This would allow LC15 to respond to faster velocity OFF bars and motion-defined stimuli.

Activation of mALB4 would turn down the inhibitory tone provided by the GABAergic lateral neurons innervating the LC15 glomerulus (Figure S5A). mALB4 receives input from ascending AN-GNG neurons in the SEZ but these particular ANs have not been physiologically examined. A previous report demonstrated that sucrose stimulation of proboscis gustatory neurons activated the dense and diffuse arborizations of the SEZ 5-HT neurons (95). An ascending gustatory sugar signal from the SEZ 5-HT neurons could activate mALB4 and release LC15 from inhibition to help facilitate multisensory integration. Previous studies have implicated 5-HT in the modulation of feeding behavior so this ascending signal from the gustatory center may help to orchestrate state-dependent behavior (96,97).

### Serotonergic motor control and behavioral implications

Serotonin is known to mediate walking in *Drosophila*: 5-HTergic neurons in the VNC slows walking speed, whereas inhibition of those neurons increases it. 5-HT^VNC^ neurons are active during locomotion, positively correlated to the animal’s velocity. A previous report tested null receptor mutants and implicated 5-HT7 as the receptor responsible (73). Additionally, it has been reported that serotonergic cells in the VNC mediate immobility and the immobility locomotor behavior is mediated by 5-HT7 (98). If an ascending locomotor ‘stop’ signal from the GNG activates mALB4, this could release LC15 from inhibition coordinated synchronously with the volumetric release of 5-HT from 5-HTPLP. The key effects include enhancing and tuning the response gain to visual objects (Figure 6) over speeds that are relevant both for social interactions during walking and for landscape feature orientation during flight (42,49,99). By contrast, AN-GNG89 neurons could be active during locomotion and, through mALB4, release LC15 from inhibition in the absence of extracellular 5-HT. Locomotion could activate descending neurons like DNp27 that would activate LmA2, providing inhibitory tone to LC15 at the lobula. At cessation, the mALB4 neurons are no longer activated by ANs, and the PVLP101b and PVLP103 neurons are free to inhibit LC15 at the glomerulus. 5-HTPLP01 neurons are released from suppression and release 5-HT into the PVLP while 5-HT release is reduced elsewhere in the brain. 5-HTPLP01 is strongly innervated by a glutamatergic partner, PVLP106, which shows reciprocal connections and could inhibit 5-HTPLP01 from releasing 5-HT during locomotion but similarly inhibited by glutamate when 5-HTPLP01 is active (Figure S3C). This increase in inhibitory tone could sharpen the features in the visual scene upon cessation of locomotion.

Serotonin acts as both a synaptic neurotransmitter and an extrinsic neuromodulator in invertebrate and vertebrate systems. 5-HT acts via ionotropic signaling by binding to and opening ion channels in the postsynaptic sites of target cells (100). Additionally, 5-HT acts upon metabotropically by binding to receptors in the cell membranes of a local extracellular space, activating GPCR cascades in multiple heterogeneous cell types. There is a large body of literature on functional mechanisms of serotonergic neurotransmission for both vertebrates and invertebrates, but the neurohormonal mechanisms of 5-HT are critically understudied. The flexibility that 5-HT provides to sensory and motor circuits and the overall contribution of paracrine signaling to serotonergic signaling is unknown.

In both cases of synaptic and paracrine signaling, we know that 5-HT shifts the excitation-inhibition relationship at critical behavioral moments. Inhibitory regulation of circuit elements by 5-HT is not exclusive to fruit flies. In locusts, 5-HT is responsible for the state change from the solitary to gregarious swarming phase, triggering phenotypic, behavioral, and physiological changes in the object-detecting descending contralateral motion detector (DCMD) (17,101). In crabs, upon eliminating this input to the A1 cells, the spontaneous firing rates of 5-HT cells are increased by about 50%, suggesting that these cells are under constant inhibitory regulation (102,103). In crayfish, pressure injection of 5-HT causes spatially-segregated inhibitory effects synaptically and excitatory effects through paracrine action on a motor neuron (104). In the visual cortex of mice, it was observed that visual stimulation with a light flash triggers excitatory and inhibitory conductances that are staggered by a few milliseconds (105). Upon visual feature presentation, the ratio between excitation and inhibition is initially tilted toward excitation, and subsequently shifts toward inhibition. The rapid change in the excitation-inhibition ratio plays a critical role in spatial and temporal response tuning. It has previously been demonstrated in vertebrates that oriented OFF bars in the visual field (106,107) lead to the concomitant occurrence of synaptic excitation and inhibition in sensory cortices (also see (108). Future research on the distribution of 5-HT receptors, as well as the function of each receptor on each LC, would be required to resolve these outstanding issues and propel our understanding of mechanisms of neuromodulatory action on visual processing.

## Glossary

5-HT: Serotonin, 5-hydroxytryptamine
5-HTR: Serotonin receptor
AVLP: Anterior ventrolateral protocerebrum
CPG: Central pattern generator
GABA: γ-aminobutyric acid
GNG: Gnathal ganglion
GPCR: G protein-coupled receptor La Lamina
LC: Lobula columnar cells
Lo: Lobula
LN: Lateral neuron
LP: Lobula plate
Me: Medulla
OG: Optic glomerulus
ORN: Olfactory receptor neuron
PVLP: Posterior ventrolateral protocerebrum
PN: Projection neurons
SEZ: Subesophageal zone
Trh: Tryptophan hydroxylase
VPN: Visual projection neuron

## Data availability

Source code and other files supporting this project can be found at https://github.com/DJBertsch/LC15-serotonin. Original datasets for this study are available upon request.

## Supplemental material

Supplemental Fig. S1: [https://doi.org/10.5281/zenodo.15947777] Supplemental Fig. S2: [https://doi.org/10.5281/zenodo.15948201] Supplemental Fig. S3: [https://doi.org/10.5281/zenodo.15948332] Supplemental Fig. S4: [https://doi.org/10.5281/zenodo.15948405] Supplemental Fig. S5: [https://doi.org/10.5281/zenodo.15948459] Supplemental Fig. S6: [https://doi.org/10.5281/zenodo.15948493]

## Acknowledgements

The authors thank SL Bonanno and DE Krantz for helpful discussions and reagents. We also thank the anonymous reviewers for their insightful comments.

## Author contributions

DJB designed the experiments, performed imaging experiments, wrote analysis code, analyzed data, interpreted results, prepared figures, and wrote the manuscript. LMPC provided genetics assistance and edited the manuscript. MAF acquired funding, conceived of the scope of work, designed experiments, provided supervision, and wrote the manuscript.

## Supplemental Figures

**Figure S1.**
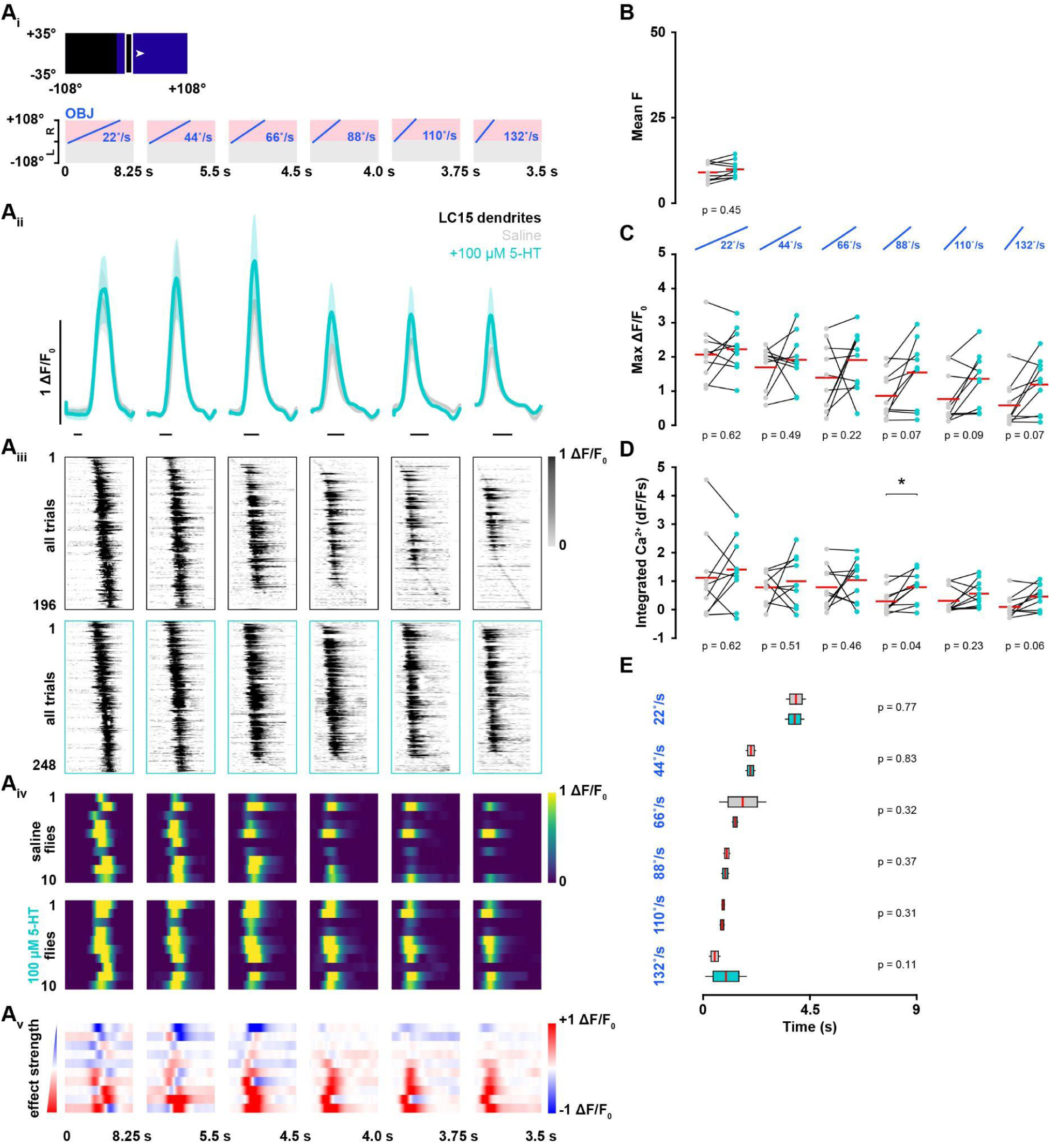
5-HT recruits individual LC15 dendrites in response to high velocity OFF bars. (Ai) Schematic illustrating the 8.8° x 70° vertical bar presented on the LED display (top) and the motion of the translating feature over time (bottom). The OFF bar moved at six different velocities (22, 44, 66, 88, 110, and 132 °/s) across the right side of the LED display. (Aii) Population mean GCaMP6f signal from the LC15 dendrites (± SEM shading) before (gray) and after 100 µM 5-HT (cyan). The vertical scale bar (black) represents 1 ΔF/F_0_. The horizontal scale bar (black) represents 1 second. (Aiii) ΔF/F_0_ response raster plots displaying individual trial responses from all ROIs. Two repeats of each stimulus were shown to each animal before (N = 10, n = 196) and after 100 µM 5-HT (n = 248). The trial responses are sorted by the time of the peak ΔF/F_0_ response. (Aiv) ΔF/F_0_ response raster plots displaying mean animal responses before (top) and after 100 µM 5-HT (bottom) (N = 10). Flies 1-10 (rows) are sorted the same in the pre-5-HT (top) and post-5-HT (bottom) condition rasters. The bottom row of each raster shows the population mean, denoted with Σ. The vertical red line represents the mean time of the ΔF/F_0_ peaks, the box indicates the peak of the ΔF/F_0_ mean trace. (Av) Mean change in effect size rasters of LC15 dendrites GCaMP6f responses for each animal in the saline to the 5-HT experimental group. Animals (row) are sorted by increasing effect size strength. The bottom row of each raster shows the change in the population mean, denoted with Σ. The vertical red line represents the mean time of the ΔF/F_0_ peaks, and the vertical blue line represents the mean time of the ΔF/F_0_ troughs.The red box indicates the peak of the ΔF/F_0_ mean trace and the blue box indicates the trough of the ΔF/F_0_ mean trace. (B) Pairwise comparison of mean fluorescent intensity in the LC15 glomerulus before (gray) and after 100 µM 5-HT (cyan). Horizontal red lines correspond to the population means. Two-tailed paired t-test, N = 10. Saline mean F = 9.3 ±2.7F. 5-HT mean F = 10.2 ±2.4F. *p* = 0.45951. Mean time lapse between pre- and post-conditions = +40 minutes. (C) Pairwise comparison of maximum ΔF/F_0_ responses in the LC15 glomerulus before (gray) and after 100 µM 5-HT (cyan). Two-tailed paired t-test, N = 10. Saline mean responses: 2.08 ±0.75, 1.70 ±0.64, 1.40 ±0.97, 0.87 ±0.67, 0.77 ±0.74, and 0.59 ±0.64 ΔF/F_0_. 5-HT mean responses: 2.23 ±0.64, 1.92 ±0.76, 1.92 ±0.88, 1.55 ±0.95, 1.36 ±0.74, and 1.20 ±0.78 ΔF/F_0_. *p* = 0.62493; *p* = 0.49094; *p* = 0.22677; *p* = 0.079596; *p* = 0.099429; *p* = 0.070617. (D) Pairwise comparison of integrated Ca^2+^ responses in the LC15 glomerulus before (gray) and after 100 µM 5-HT (cyan). Two-tailed paired t-test, N = 10. Saline mean responses: 1.16 ±1.49, 0.82 ±0.54, 0.82 ±0.75, 0.33 ±0.45, 0.35 ±0.46, and 0.13 ±0.40. 5-HT mean responses: 1.45 ±1.12, 1.04 ±0.88, 1.07 ±0.74, 0.83 ±0.56, 0.60 ±0.46, and 0.49 ±0.44. *p* = 0.62567; *p* = 0.51522; *p* = 0.46189; *p* = 0.044255*; *p* = 0.23225; *p* = 0.069583. (E) Pairwise comparison of time to half-max (s) for LC15 glomerulus responses before (gray) and after 100 µM 5-HT (cyan). Two-tailed paired t-test, N = 10. Saline mean responses: 3.9 ±0.4, 2.0 ±0.2, 1.7 ±1.0, 1.0 ±0.2, 0.9 ±0.1, and 0.5 ±0.3 seconds. 5-HT mean responses: 3.9 ±0.4, 2.0 ±0.2, 1.4 ±0.2, 0.9 ±0.2, 0.8 ±0.1, and 1.0 ±0.9 seconds. *p* = 0.77929; *p* = 0.83323; *p* = 0.3271; *p* = 0.37659; *p* = 0.31123; *p* = 0.11728.

**Figure S2.**
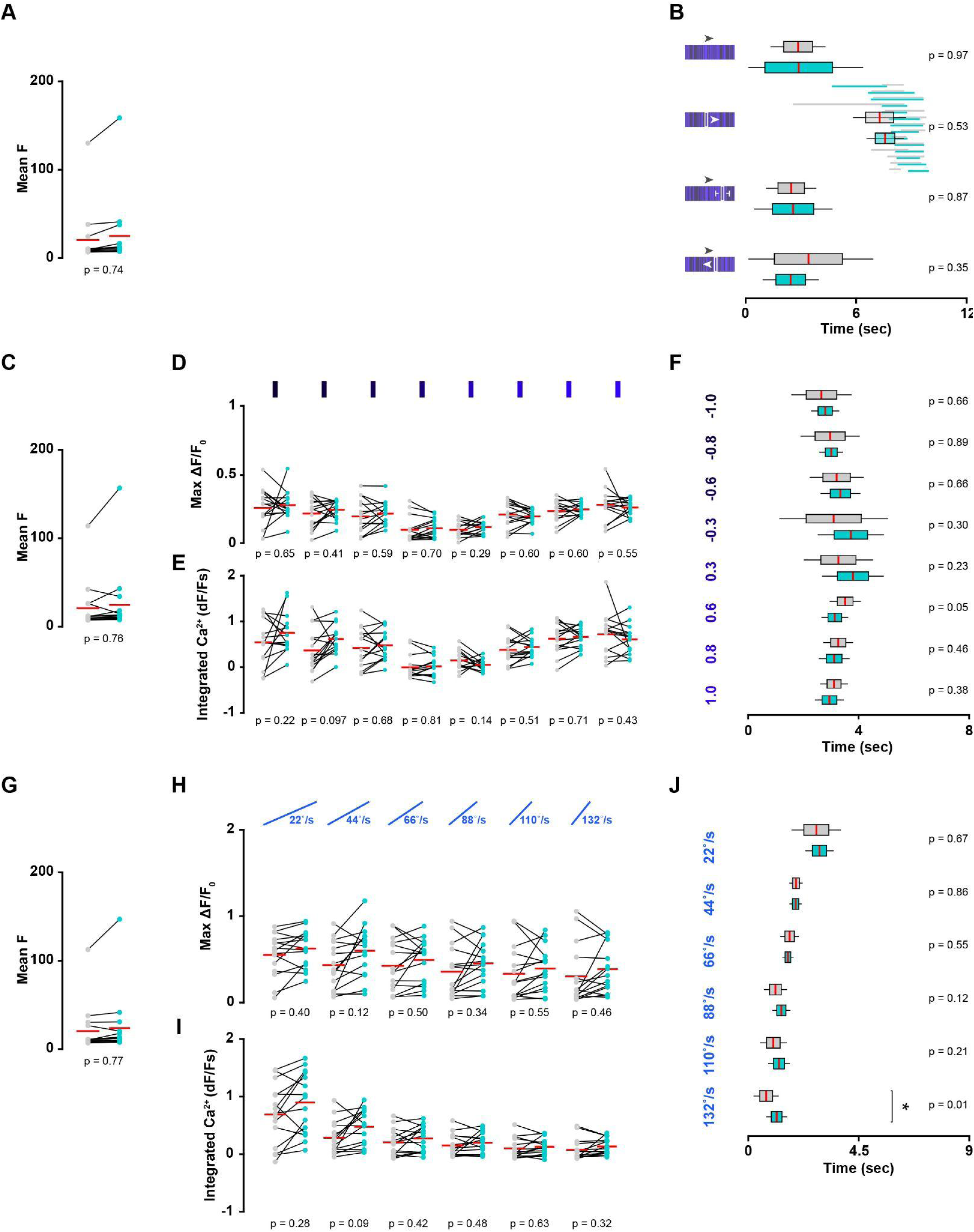
Accompanying statistics for Figure 5. (A) Pairwise comparison of mean fluorescent intensity in the LC12 glomerulus before (gray) and after 100 µM 5-HT (cyan). Horizontal red lines correspond to the population means. Two-tailed paired t-test, N = 14. Saline mean F = 20.5 ± 32.7F. 5-HT mean F = 25.1 ± 39.9F. *p* = 0.74341. Mean time lapse between pre- and post-conditions = +39 minutes. (B) Pairwise comparison of time to half-max (s) for LC12 glomerular responses before (gray) and after 100 µM 5-HT (cyan). Two-tailed paired t-test, N = 14. Saline mean responses: 2.7 ±1.5, 7.1 ±1.5, 2.3 ±1.4, and 3.3 ±3.5 seconds. 5-HT mean responses: 2.7 ± 3.5, 7.4 ± 1.0, 2.4 ± 2.1, and 2.3 ±1.5 seconds. *p* = 0.97536; *p* = 0.53507; *p* = 0.87825; *p* = 0.35728. The horizontal lines behind the box plots in the motion-defined bar condition indicate the time period from the half-max to the maximum ΔF/F_0_ responses for each animal before (gray) and after 100 µM 5-HT (cyan). (C) Pairwise comparison of mean fluorescent intensity in the LC12 dendrites before (gray) and after 100 µM 5-HT (cyan). Horizontal red lines correspond to the population means. Two-tailed paired t-test, N = 15. Saline mean F = 20.9 ±27.6F. 5-HT mean F = 24.6 ±37.9F. *p* = 0.76034. Mean time lapse between pre- and post-conditions = +38 minutes. (D) Pairwise comparison of maximum ΔF/F_0_ responses in the LC12 glomerulus before (gray) and after 100 µM 5-HT (cyan). Two-tailed paired t-test, N = 15. Saline mean responses: 0.26 ±0.14, 0.22 ±0.11, 0.20 ±0.12, 0.10 ±0.09, 0.10 ±0.05, 0.21 ± 0.09, 0.24 ±0.08, and 0.28 ±0.11 ΔF/F_0_. 5-HT mean responses: 0.28 ±0.10, 0.25 ±0.07, 0.22 ±0.08, 0.11 ±0.07, 0.12 ±0.05, 0.20 ±0.05, 0.25 ±0.04, and 0.26 ±0.06 ΔF/F_0_. *p* = 0.65388; *p* = 0.419; *p* = 0.59623; *p* = 0.70333; *p* = 0.29775; *p* = 0.6008; *p* = 0.60734; *p* = 0.55675. (E) Pairwise comparison of integrated Ca^2+^ responses in the LC12 glomerulus before (gray) and after 100 µM 5-HT (cyan). Two-tailed paired t-test, N = 15. Saline mean responses: 0.54 ±0.51, 0.37 ±0.47, 0.42 ±0.44, -0.01 ±0.26, 0.15 ±0.21, 0.38 ±0.30, 0.62 ±0.34, and 0.72 ±0.46. 5-HT mean responses: 0.75 ±0.43, 0.62 ±0.32, 0.48 ±0.33, 0.02 ±0.21, 0.05 ±0.13, 0.44 ±0.20, 0.66 ±0.21, and 0.61 ±0.32. *p* = 0.22942; *p* = 0.097839; *p* = 0.68063; *p* = 0.81441; *p* = 0.14218; *p* = 0.51476; *p* = 0.71213; *p* = 0.43841. (F) Pairwise comparison of time to half-max (s) for LC12 glomerulus responses before (gray) and after 100 µM 5-HT (cyan). Two-tailed paired t-test, N = 15. Saline mean responses: 2.65 ±1.09, 2.96 ±1.07, 3.20 ±0.98, 3.09 ±1.97, 3.26 ±1.26, 3.51 ±0.56, 3.26 ±0.54, and 3.10 ±0.50 seconds. 5-HT mean responses: 2.78 ±0.51, 3.00 ±0.43, 3.34 ±0.72, 3.71 ±1.20, 3.79 ±1.12, 3.13 ±0.49, 3.11 ±0.56, and 2.94 ±0.53 seconds. *p* = 0.66706; *p* = 0.89636; *p* = 0.66549; *p* = 0.30449; *p* = 0.23448; *p* = 0.058331; *p* = 0.46895; *p* = 0.38026. (G) Pairwise comparison of mean fluorescent intensity in the LC12 glomerulus before (gray) and after 100 µM 5-HT (cyan). Two-tailed paired t-test, N = 15. Saline mean F = 20.4 ±27.2F. 5-HT mean F = 23.8 ±35.4F. *p* = 0.77095. Mean time lapse between pre- and post-conditions = +38 minutes. (H) Pairwise comparison of maximum ΔF/F_0_ responses in the LC12 glomerulus before (gray) and after 100 µM 5-HT (cyan). Two-tailed paired t-test, N = 15. Saline mean responses: 0.28 ±0.12, 0.22 ±0.14, 0.21 ±0.15, 0.18 ±0.15, 0.17 ±0.16, and 0.15 ±0.17 ΔF/F_0_. 5-HT mean responses: 0.31 ±0.11, 0.30 ±0.15, 0.25 ±0.13, 0.23 ±0.13, 0.20 ±0.13, and 0.19 ±0.14 ΔF/F_0_. *p* = 0.40072; *p* = 0.1233; *p* = 0.50841; *p* = 0.34823; *p* = 0.55249; *p* = 0.46837. (I) Pairwise comparison of integrated Ca^2+^ responses in the LC12 glomerulus before (gray) and after 100 µM 5-HT (cyan). Two-tailed paired t-test, N = 15. Saline mean responses: 0.69 ±0.53, 0.28 ±0.29, 0.21 ±0.23, 0.15 ±0.19, 0.10 ±0.18, and 0.07 ±0.19. 5-HT mean responses: 0.90 ±0.53, 0.47 ±0.31, 0.27 ±0.22, 0.20 ±0.18, 0.13 ±0.15, and 0.13 ±0.14. *p* = 0.28879; *p* = 0.090397; *p* = 0.42695; *p* = 0.4842; *p* = 0.63555; *p* = 0.32845. (J) Pairwise comparison of time to half-max (s) for LC12 glomerulus responses before (gray) and after 100 µM 5-HT (cyan). Two-tailed paired t-test, N = 15. Saline mean responses: 2.8 ±1.0, 1.9 ±0.3, 1.7 ±0.4, 1.1 ±0.5, 1.0 ±0.6, and 0.7 ±0.5 seconds. 5-HT mean responses: 2.9 ±0.6, 1.9 ±0.3, 1.6 ±0.2, 1.4 ±0.4, 1.3 ±0.4, and 1.2 ±0.4 seconds. *p* = 0.67116; *p* = 0.86005; *p* = 0.55694; *p* = 0.12728; *p* = 0.21319; *p* = 0.019491*.

**Figure S3.**
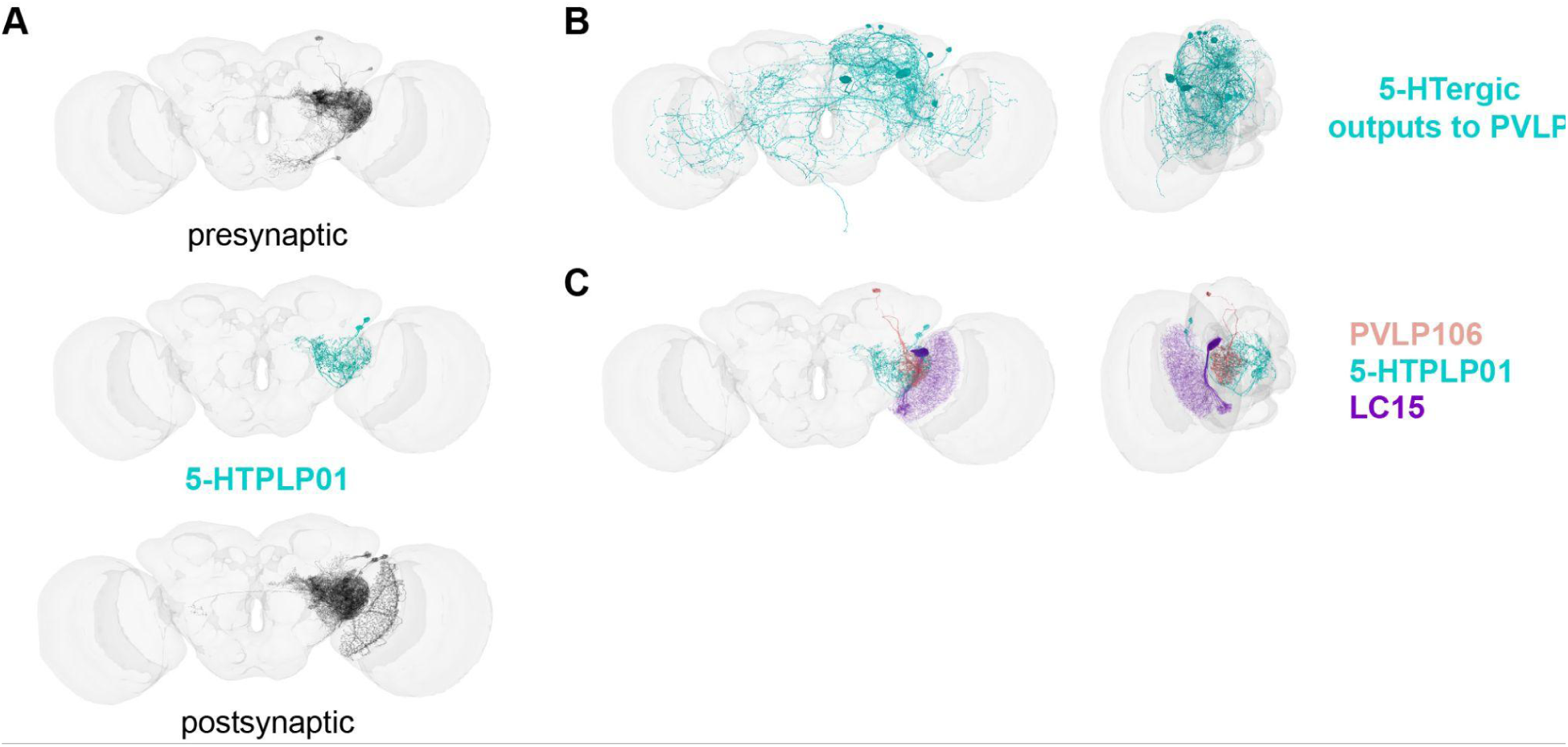
Serotonergic innervation of the PVLP. (A) Presynaptic partners (black, top) to 5-HTPLP (cyan) and postsynaptic partners (black, bottom) in the right hemisphere. (B) Serotonergic neurons (cyan) that have output synapses in the right hemisphere PVLP. (C) Morphology of PVLP106, a glutamatergic interneuron that has reciprocal synapses onto 5-HTPLP01 (cyan). LC15 (purple) does not make synapses with either group.

**Figure S4.**
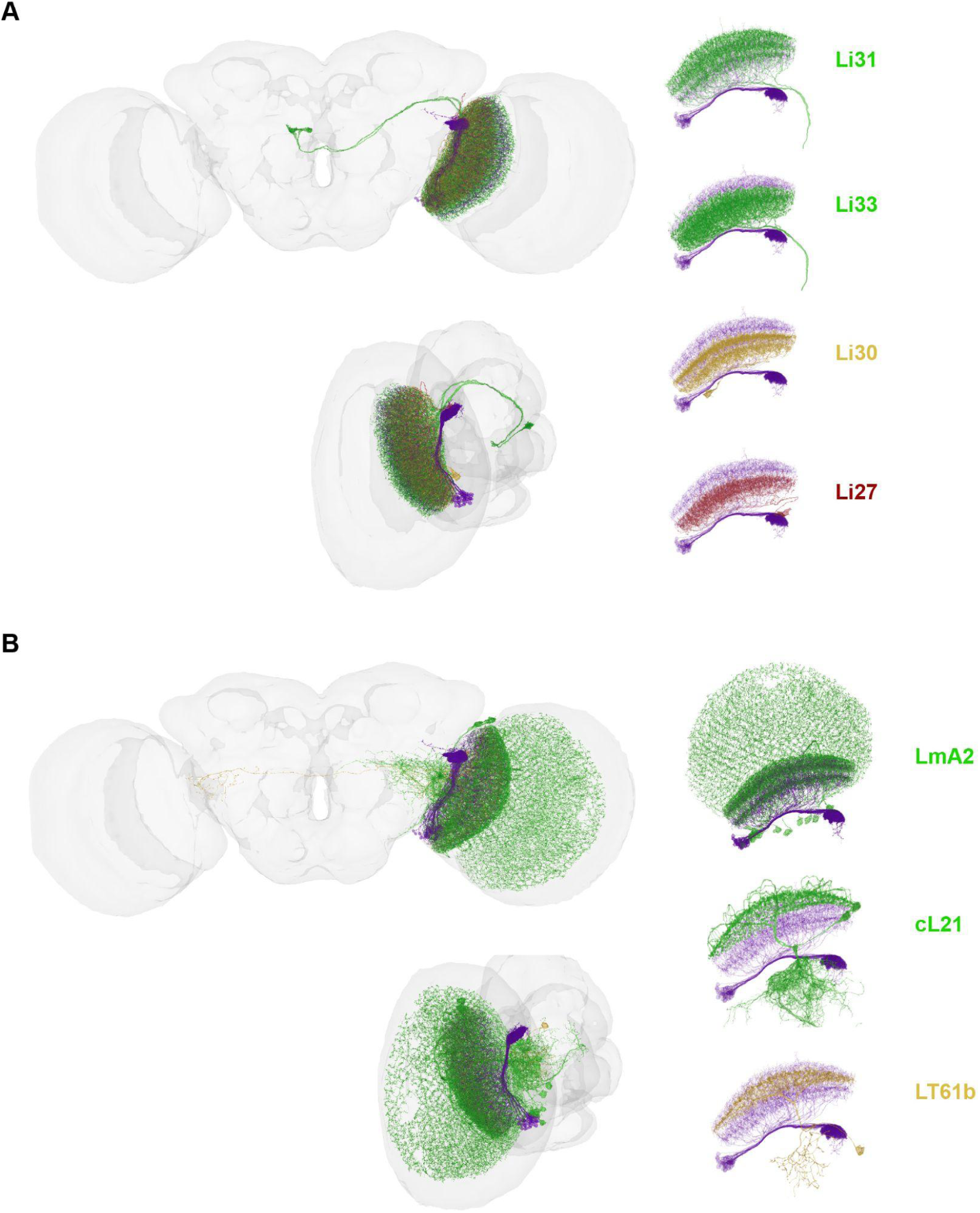
LC15 inhibitory circuit motifs in the optic lobe. (A) Morphology reconstruction of Li30-Li31-Li33-Li27 circuit motif. LC15 (purple) is shown for reference. The neurons are colored according to neurotransmitter type (yellow, cholinergic) (green, GABAergic) (red, glutamatergic). (B) Morphology reconstruction of LT61b-cL21-LmA2 circuit motif. LC15 (purple) is shown for reference. The neurons are colored according to neurotransmitter type (yellow, cholinergic) (green, GABAergic). The centrifugal GABAergic neurons, cL21, is the highest downstream partner of 5-HTPLP01, making over 700 glutamatergic synapses. In the lobula, inhibition of cL21 would release LmA2 neurons from GABAergic inhibition, causing a stronger inhibitory suppression on LC15 at the lobula. Glutamatergic suppression of cL21 would also release T2 and T3 from inhibition at the lobula, possibly causing a potentiation of small objects in other LCs. Opposingly, LT61b, a cholinergic output of LC15, sends excitatory signals from the VLP on both hemispheres to cL21 in the lobula. Activation of cL21 by LT61b would strengthen the GABAergic inhibition of cL21 over LmA2 and release LC15‘s dendrites from inhibition.

**Figure S5.**
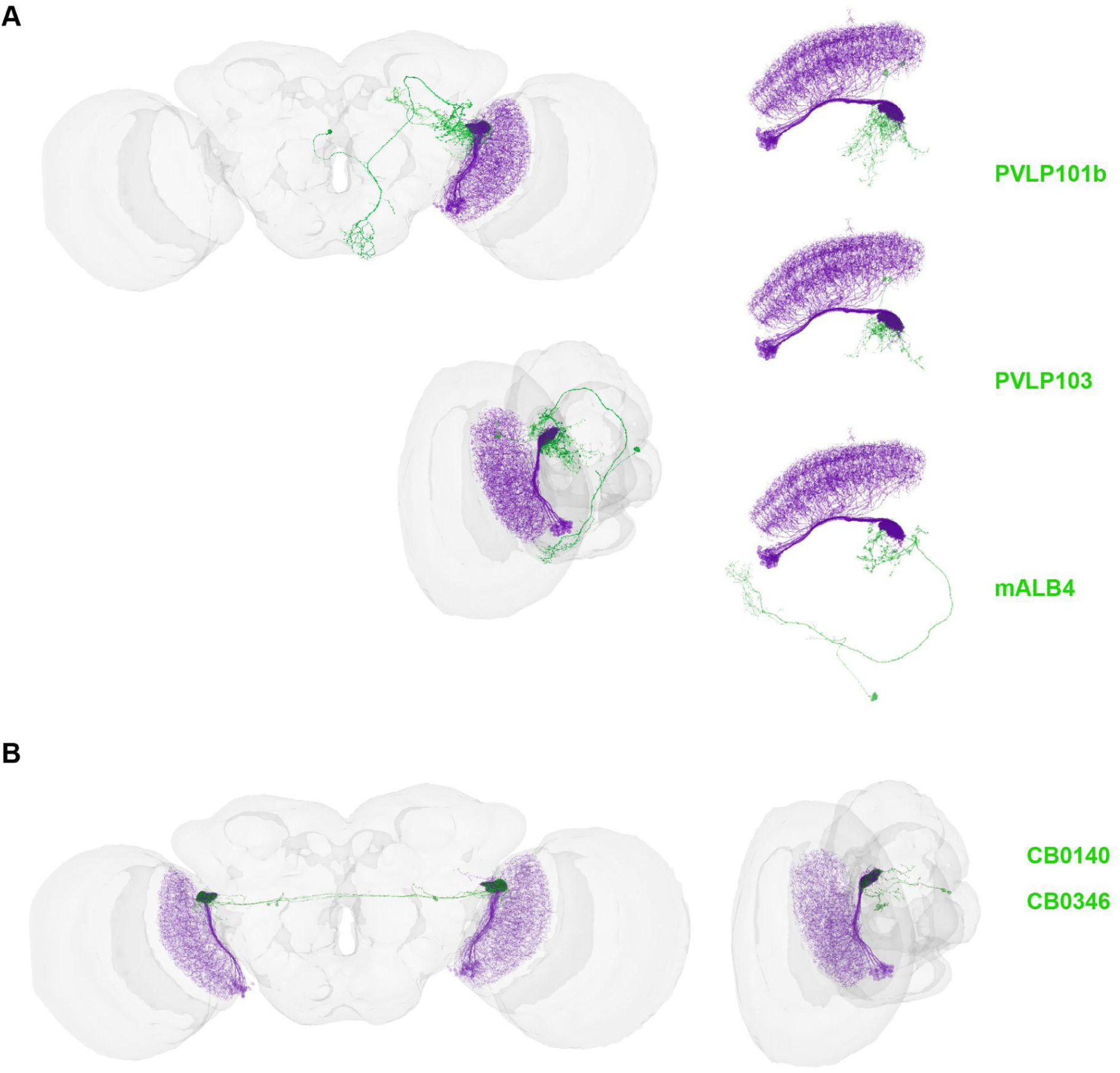
LC15 inhibitory circuit motifs in the central brain. (A) Morphology reconstruction of mALB4-PVLP101b-PVLP103 circuit motif. LC15 (purple) is shown for reference. The neurons are colored according to neurotransmitter type (green, GABAergic). (B) Morphology reconstruction of CB0140-CB0346 circuit motif. LC15 (purple) is shown for reference. The neurons are colored according to neurotransmitter type (green, GABAergic).

**Figure S6.**
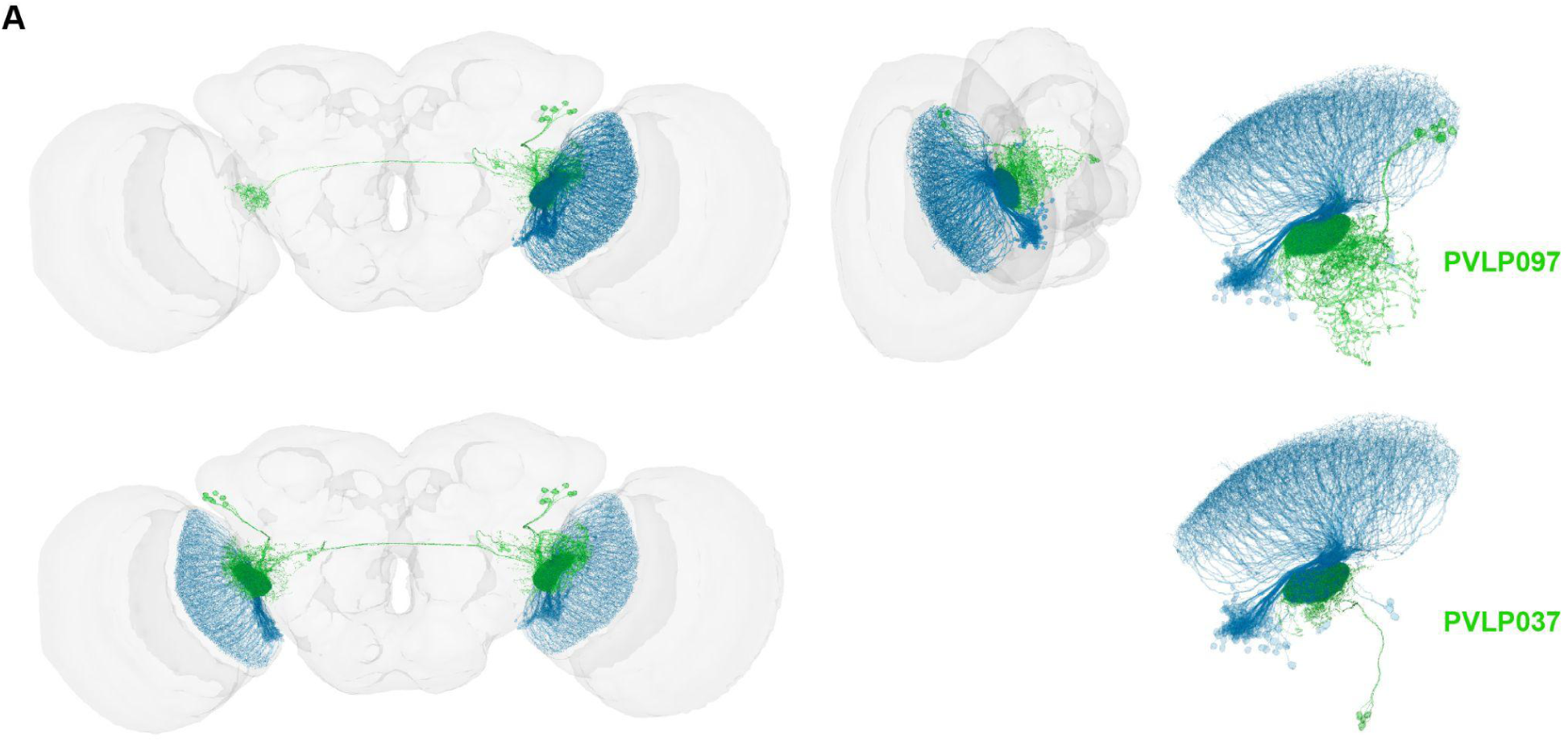
LC12 axo-axonal inhibitory circuit motif. (A) Morphology reconstruction of PVLP037-PVLP097 circuit motif. LC12 (blue) is shown for reference in one hemisphere (left, top), and bilaterally (left, bottom). The neurons are colored according to neurotransmitter type (green, GABAergic). This circuit motif is reminiscent of what has previously been reported in other LCs (59,109). There are 7 PVLP097 cells that make over 2,000 GABAergic synapses back onto LC12. High inhibitory tone from PVLP097 likely contributes to ‘weak’ glomerular responses from LC12 compared to previous reports that imaged in the lobula dendrites. Instead of direct axo-axonic LC12-LC12 connections by a GABAergic lateral neuron, the PVLP037 cells make 684 GABAergic synapses onto PVLP097. If PVLP037 is a state-dependent neuron, it could mediate a switch for context-specific LC12 feature sensitivity. For example, flight could activate PVLP037 and release LC12 from inhibition to better respond to a vertical bar, or looming stimulus. Removing the PVLP097 cells should expand the feature-detection parameter space in LC12 and activating PVLP037 may recapitulate that effect.

